# Neurodevelopmental Subtypes of Functional Brain Organization in the ABCD Study Using a Rigorous Analytic Framework

**DOI:** 10.1101/2024.03.16.585343

**Authors:** Jacob DeRosa, Naomi P. Friedman, Vince Calhoun, Marie T. Banich

## Abstract

The current study demonstrates that an individual’s resting-state functional connectivity (RSFC) is a dependable biomarker for identifying differential patterns of cognitive and emotional functioning during late childhood. Using baseline RSFC data from the Adolescent Brain Cognitive Development (ABCD) study, which includes children aged 9-11, we identified four distinct RSFC subtypes We introduce an integrated methodological pipeline for testing the reliability and importance of these subtypes. In the Identification phase, Leiden Community Detection defined RSFC subtypes, with their reproducibility confirmed through a split-sample technique in the Validation stage. The Evaluation phase showed that distinct cognitive and mental health profiles are associated with each subtype, with the Predictive phase indicating that subtypes better predict various cognitive and mental health characteristics than individual RSFC connections. The Replication stage employed bootstrapping and down-sampling methods to substantiate the reproducibility of these subtypes further. This work allows future explorations of developmental trajectories of these RSFC subtypes.

## 1. Introduction

The study of neurodevelopment is essential for elucidating the intricate processes and mechanisms underpinning children’s cognitive abilities, emotions, and social growth. The Adolescent Brain Cognitive Development (ABCD) study is an extensive longitudinal effort to identify the underlying relationships among biological, environmental, and social factors influencing brain development and cognitive functioning during late childhood and adolescence (Volkow et al., 2018). The ABCD study is designed to identify critical determinants of substance use, mental health, and cognitive functioning, all important facets of adolescent development (Casey et al., 2018). However, achieving accurate predictions of brain-behavior relationships remains a considerable challenge (Rosenberg et al., 2018). A possible obstacle is the inherent heterogeneity of neurodevelopment, which does not follow similar patterns across all children. Machine learning-based subtyping methods, such as clustering, have gained traction to address challenges associated with assessing heterogeneity in neurodevelopment by identifying distinct profiles and revealing associations with cognitive functioning and characteristics linked to psychopathology (DeRosa, Rosch, et al., 2023; Gupta et al., 2017; Nikolaidis et al., 2022). If different subgroups of children/adolescents have distinct brain profiles, pronounced variability in brain-behavior relationships may be observed (Bathelt et al., 2018; Crone & Elzinga, 2015).

Identifying different subgroups in neurodevelopmental studies exemplifies the concept of nested heterogeneity, which can be studied through the lens of precision medicine and individualized treatment strategies (DeRosa, Rosch, et al., 2023; Fair et al., 2012; Fekson et al., 2023). This approach acknowledges the diverse pathways of brain development and the variation in cognitive and mental health outcomes, aiming to tailor interventions and understandings to the unique profiles of individuals. In contrast to non-nested heterogeneity, nested heterogeneity refers to the presence of multiple layers of variability within a system. While non-nested heterogeneity implies a singular level of diversity, nested heterogeneity indicates that further distinct variations exist within each subgroup or category. In neurodevelopment, nested heterogeneity underscores that variations in brain development are not uniform; instead, they manifest across multiple levels (the nested layers). These levels range from individual differences to distinct subgroup characteristics shaped by biological, environmental, and social factors.

Given nested heterogeneity’s potential contribution to the limitations in predicting brain-behavior relationships, researchers are increasingly interested in exploring it in the field of child development (Feczko & Fair, 2020). A thorough analysis of the intricate variations in neurodevelopmental profiles within the ABCD dataset provides an opportunity for improving our understanding and predictive ability concerning the complex relationship between brain development and behavior. The current paper aims to provide an empirical investigation and framework of the degree to which identifying subgroups of individuals based on their pattern of resting-state functional connectivity (RSFC) at the initial time point of the ABCD study might aid in revealing brain-behavior relationships. The ABCD study offers an unprecedented dataset of over 11,000 individuals on whom brain measures of anatomy (grey and white matter) and functional activation (resting-state, task-based) have been obtained, along with multiple behavioral measures of cognitive and emotional function. As such, it is an ideal sample to pursue this issue.

### 1.1. Resting State Functional Connectivity

We propose leveraging RSFC to address the inherent heterogeneity in connectivity across brain networks. Numerous previous studies have emphasized the usefulness of RSFC in identifying connectivity patterns between brain regions (Cohen et al., 2008; Dosenbach et al., 2007; Fair et al., 2009; Power et al., 2011; Yeo et al., 2011). A compelling resemblance between resting-state and task-evoked networks has been observed, indicating a tight relationship between the brain’s intrinsic network architecture in a resting state and its functional organization during task execution (Cole et al., 2014). Notably, RSFC yields consistent stability over time and maintains its robustness irrespective of changes in task performance or state for a given individual, establishing it as a dependable form of functional neuroimaging data for longitudinal research on individual differences (Reineberg & Banich, 2016). This characteristic of RSFC, paired with the large-consortia data from ABCD, provides an excellent dataset to extract RSFC subtype profiles. Furthermore, applying multivariate methodologies to the ABCD RSFC data has demonstrated that some aspects of RSFC connectivity are associated with cognitive abilities (Byington et al., 2023; Pat et al., 2022).

### 1.2. Whole-Brain Profiles as Neuro-markers of Cognitive-Emotional Subtypes

The current work is motivated by the proposition that an individual’s whole-brain RSFC profile can be a reliable neuro-marker for characterizing distinct development patterns. We aim to identify RSFC subtypes that may help elucidate the spectrum of cognitive functioning and mental health patterns during late childhood. This endeavor seeks to reveal nested heterogeneity in the ABCD baseline sample, where multiple unique functional connectivity patterns may coexist to varying degrees within the same population, each distinctly associated with various cognitive functioning and mental health outcomes (Ohashi & Ostry, 2021; Peverill et al., 2019). Nested heterogeneity here would be characterized not by a hierarchical structure but by the coexistence of diverse, unique functional connectivity patterns within the baseline sample. Each RSFC subtype would then be assessed to evaluate if it is linked to specific demographic, cognitive, and mental health indicators.

Notably, these RSFC subtypes differ from “neuroimaging fingerprinting” analyses, which identify unique individual-level patterns of brain connectivity (Finn et al., 2015). These RSFC subtype profiles enable the characterization and comparison of distinct connectivity patterns, providing a valuable method to categorize and analyze common connectivity patterns at the (sub)group level. They capture the heterogeneity of RSFC within a population and offer insights into the diversity of functional brain organization and its relationship to individual differences in experiences or traits. Therefore, rather than replacing individual-level analyses, such as fingerprinting, these subtypes complement and enhance our understanding of individual differences in brain organization (Fu, Liu, et al., 2022; Fu, Sui, et al., 2022).

### 1.3. The IVEPR Framework: A Standardized Subtyping Evaluation Approach

To optimally identify subtypes based on RSFC profiles, it is critical to ensure the robustness and reliability of such subtypes. Such assurance is crucial if these subtypes are to be regarded as meaningful. To maximize the rigor of the present work, we introduce the Identification, Validation, Evaluation, Prediction, and Replication (IVEPR) framework (**Figure 1** contains a detailed outline of the steps in this framework). The IVEPR framework is a strategic combination of existing methodologies (Byington et al., 2023; DeRosa, Rosch, et al., 2023; Nikolaidis et al., 2021; Pat et al., 2022, 2023) that provides a cohesive, standardized procedure for rigorous subtype identification. This unique integration fosters reliable identification and validation of the RSFC subtypes, providing a solid foundation for assessing their utilization as neuro-markers. The framework’s comprehensive design addresses robustness and reproducibility issues commonly seen in data-driven clustering research (Arbabshirani et al., 2017; Bzdok, 2017; Demirci et al., 2008; Varoquaux et al., 2017) thereby enhancing the credibility of the findings. The IVEPR framework sets a high standard for analytical precision that robustly evaluates and validates the RSFC subtypes, bolstering our ability to leverage these potentially impactful neuro-markers in neurodevelopmental research and applications and paving the way for their longitudinal tracking.

**Figure 1.**
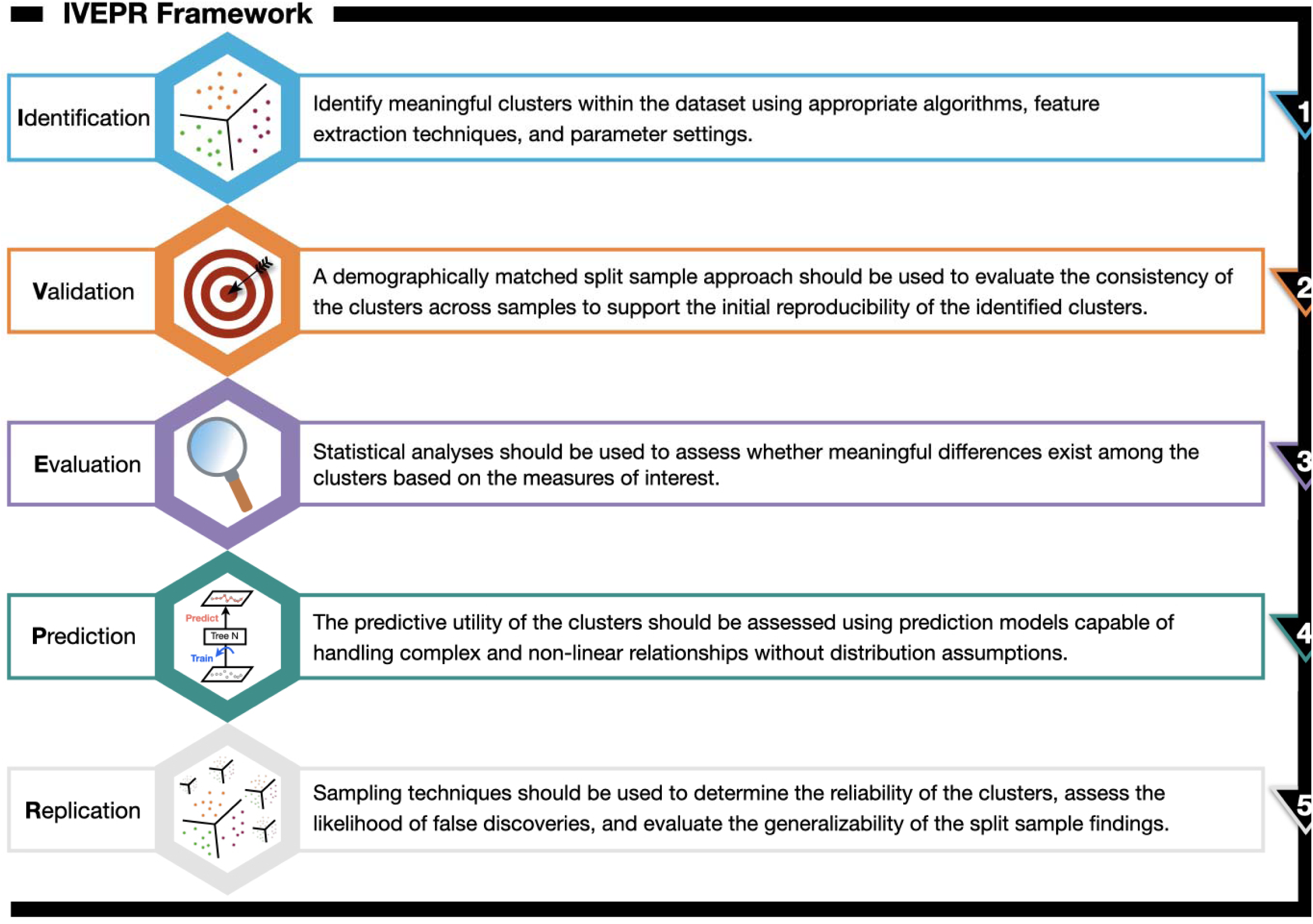
The *IVEPR* Framework - An Integrated Solution for Data-Driven Clustering in Neuroimaging Subtyping. *Note:* We use the term “subtype” to refer to the clusters of individual identified as the outputs of the data-driven Leiden Community Detection analyses.

### 1.4. Present Applications

This study identifies RSFC subtypes and then investigates their associations with cognitive functioning, mental health, and demographic attributes in a cohort of 9-11-year-old children using data from the baseline scan of the ABCD study. First, we seek to identify and validate distinct RSFC subtypes using multivariate techniques. Next, we explore whether these subtype differ in their ability to predict cognitive abilities and mental health. Finally, we evaluate the reproducibility and reliability of the identified subtypes and their predictive abilities through rigorous statistical analyses. In the context of the present report, the following terms are defined as follows: 1) Reproducibility refers to the ability to obtain the same results using the same dataset and analytical methods. Here, it means that the identified RSFC subtypes and their behavioral differences and predictive abilities for cognitive abilities and mental health should be able to be re-identified or re-predicted across two ABCD Reproducible Matched Samples (Feczko et al., 2021). 2) Reliability refers to the consistency of the results obtained from our approach; that is, the identified RSFC subtypes consistently yield similar relationships to behavior across different observations or assessments. 3) Robustness refers to the ability of the identified RSFC subtypes to remain stable and accurate across analytic variations, such as changes in sample size or composition or slight deviations in analysis methodology. **Figure 2** contains a schematic representation of our analysis pipeline.

**Figure 2.**
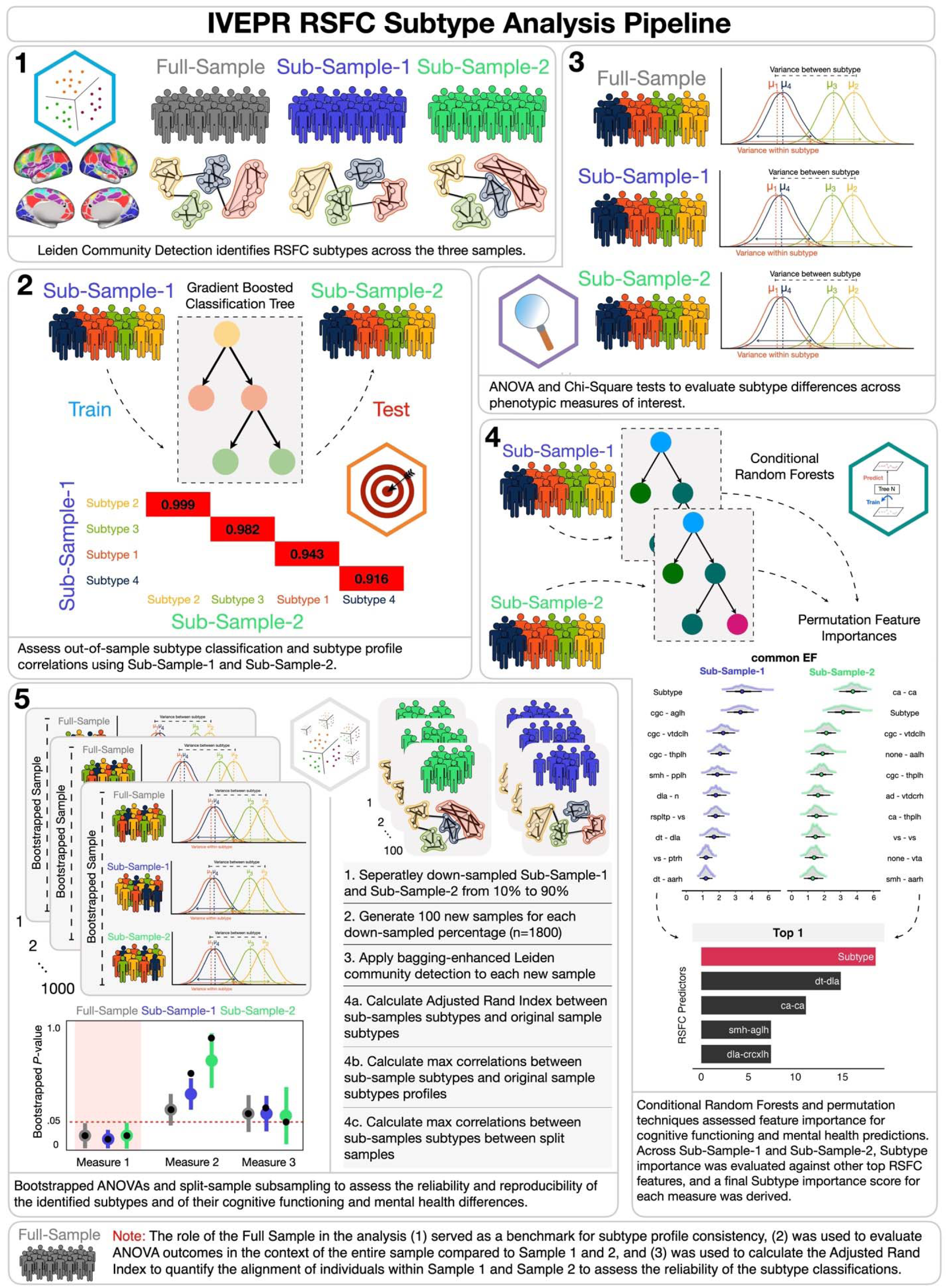
Schematic representation of the RSFC subtyping analysis pipeline steps according to the IVEPR framework. Note: The IVEPR framework icon labels are included for each stage of the analysis pipeline.

We begin by investigating whether subtyping can help to shed light on differences in cognitive and emotional profiles in our sample. Answering whether distinct subtypes exist can improve our understanding of the underlying neural mechanisms contributing to behavior (Insel & Cuthbert, 2015). To answer this question, a range of tools and techniques drawn from previous studies will be employed, such as bootstrap-enhanced Leiden community detection for data-driven subtyping (DeRosa et al., 2023), multiple group confirmatory factor analysis (CFA) for cognitive and mental health factor extraction (Freis et al., 2022) and invariance testing, gradient-boosted decision trees and SHAPley additive explanations for subtype classification and feature importance, chi-square tests and odds ratios for subtype demographic diagnostics, and split-sample validation for ensuring the reproducibility (Byington et al., 2023; Lichenstein et al., 2022), reliability, and robustness of the identified subtypes. Employing these methods allows us to identify distinct subgroups of individuals (Gordon et al., 2016; Marek et al., 2019), classify them based on their neurobiological profiles (Fernández-Delgado et al., 2014; Woo et al., 2017) and validate our findings using split-sample and down-sampling techniques. Note that evaluating subtype invariance with regards to profiles of cognitive and emotional function is necessary for a more comprehensive understanding of the underlying neurobiological relationships between brain and behavior. If invariance is established, it will support the premise that distinct neurobiological profiles are associated with distinct patterns of cognitive and mental health processing, underscoring the broad applicability of these subtypes.

In our subtyping analysis, the key objective is to evaluate whether the RSFC subtypes provide additional predictive value for cognitive abilities and mental health problems beyond the capabilities of individual RSFC connections. In this context, individual connections are defined as the singular RSFC features that constitute features of the subtype profiles. Our focus is to determine if the RSFC subtype profiles, as integrated collections of these individual connectivity features, offer a more significant predictive utility compared to the predictive capacity of any singular RSFC feature. The goal is to establish that the collective RSFC subtype profiles can offer a more detailed and comprehensive understanding of brain-behavior relationships, surpassing the insights provided by individual connectivity features alone.

Answering this question could significantly advance our understanding of the heterogeneous nature of cognitive functioning and mental health, leading to more personalized and practical approaches to enhance cognitive development and mental health in children and adolescents (Bzdok & Meyer-Lindenberg, 2018; Whitfield-Gabrieli et al., 2016). Past research, such as studies by Nikolaidis et al. (2022, 2021), suggests that in prediction models, subtypes often outperform the individual features from which they are derived. We aim to verify whether this trend is observed with the RSFC subtype profiles within the ABCD dataset. Employing a conditional random forest (CRF) (Strobl et al., 2008) approach, our analysis is specifically designed to test whether the collective profile of these subtypes emerges as a more reliable marker for cognitive functioning and mental health compared to the predictive power of each RSFC connection. We hypothesize that the subtype profiles will consistently rank as the most important features in our CRF models, demonstrating their superiority over individual RSFC connections. This focus on utilizing RSFC subtype profiles aims to address the heterogeneous nature of cognitive functioning and mental health more precisely and contributes to developing more personalized approaches in these areas.

To date, a handful of studies have investigated subtyping-based approaches using the ABCD dataset, revealing various aspects of neurodevelopment and its links to psychopathology and cognitive functioning. For instance, one study employed multiple neuroimaging modalities to identify subgroups associated with overall psychopathology, finding that certain neurobiological profiles correlated with increased psychopathology (Wang et al., 2023). Another study used latent profile analysis on task-based fMRI ROIs to uncover seven unique neurodevelopmental profiles, each associated with distinct demographic and clinical features, indicating diverse neurodevelopmental subgroups within the population (Lichenstein et al., 2022). Additionally, a study on inhibitory control established cognitive and neurobiological profiles related to reading abilities, demonstrating significant reading ability variations among groups with different default mode network connectivity patterns (Fekson et al., 2023). Moreover, research on ADHD (Sui et al., 2023) identified two distinct ADHD biotypes with implications for personalized medication therapy, emphasizing the potential of neuroimaging markers in tailored treatment approaches (Yan et al., 2023). Together, these studies highlight the ABCD dataset’s potential role in revealing neurodevelopmental profiles that may contribute to a deeper understanding of mental health, cognitive abilities, and the potential for personalized treatment strategies.

Yet, while those previous studies have identified various neurodevelopmental profiles, our study extends deeper than simply deriving the subtype profiles, but rather critically evaluates whether these subtypes offer additional value over the metrics (in our case, RSFC connections) for predicting cognitive functioning and mental health. Our current work introduces a novel and rigorous approach focused on determining the meaningfulness and predictive power of RSFC subtypes beyond the measures used to derive them. Using the IVEPR framework, our study stands out by rigorously validating the reliability and reproducibility of the RSFC subtypes. This rigor of the multifaceted IVEPR approach aims to ensure the stability and consistency of the subtype profiles and to rigorously test their predictive utility. Moreover, our study also carefully implements safeguards against biases and overfitting by utilizing split-sample resampling and down-sampling methods at all levels of analysis. This enhances the robustness of our findings, offering a quantitatively reliable measure for assessing the reproducibility and reliability of our results. This methodology not only strengthens the validity of our conclusions but also sets a prototype for future research in the field.

## 2. STAR Methods

### 2.1. Participants

The current report used the ABCD Study Curated Annual Release 4.0, comprising 3T MRI data and cognitive assessments from 11,758 children (5,631 females) aged 9-10 years at baseline. Participants were recruited from 21 locations throughout the United States (Garavan et al., 2018). The study purposively achieved demographic (White 52.2%; Black 15.1%; Hispanic 20.4%; 3.2% including Asian, American Indian/Alaska Native, Native Hawaiian, and other Pacific Islander; Multiple races 9.2%) and socioeconomic diversity (Family annual income: <$25K - 16.1%, $25K-$49K - 15.1%, $50K-$74K - 14.0%, $75K-$99K - 14.1%, $100K-$199K - 29.5%, >$200K - 11.2%) to approximate the national demographic statistics for children of the same age as determined by the American Community Survey (Heeringa & Berglund, 2020). Please refer to **Supplemental Table 1** for a detailed examination of the demographic characteristics encompassing the samples used in the current report. Ethical aspects of the ABCD study, including informed consent, confidentiality, and sharing assessment outcomes with participants, have been discussed in depth elsewhere (Clark et al., 2018).

The ABCD study provided detailed procedures for data acquisition and MRI image processing (Casey et al., 2018; Hagler et al., 2019; Yang & Jernigan, n.d.). We followed their recommended exclusion criteria based on automated and manual quality control (QC) review of each resting-state functional magnetic resonance imaging (rs-fMRI) scan listed under the abcd_imgincl01 table (Yang & Jernigan, n.d.). The ABCD Data Analysis and Informatics Core created an exclusion flag for rs-fMRI (“imgincl_rsfmri_include”) based on several criteria involving image quality, MR neurological screening, and number of repetition times. For the current report, we removed participants with an exclusion flag for rs-fMRI and randomly selected only one sibling from each family to control for familial variance. This led to a final comprehensive sample, which we refer to as the “passed RSFC quality control” sample, consisted of 7,293 children aged between 9 and 10.9 years. However, we also conducted two important supplementary subtyping analyses. The first supplementary set of analyses involved all participants with baseline RSFC data, and the second supplementary set of analyses considered only those that were designated to be excluded under the “imgincl_rsfmri_include”. Our supplementary analyses with all (randomly selected sibling) sample, which we refer to as the “complete” sample, with RSFC data comprised a sample size of 9,027. Participants with an exclusion flag, which we refer to as the “high motion” sample, formed a sample size of 1,293. Refer to **Supplemental Tables 3-4** for demographics of the “complete sample” and the “high-motion” sample.

The two split samples (Sub-Sample-1 and Sub-Sample-2) used in this study, predefined by the ABCD study, were matched on nine developmental factors (site, age, sex, ethnicity, grade, parental education, handedness, family income, and structure), plus anesthesia exposure, to consider its timing (lifespan or perinatal) and effects on behavioral and neurodevelopmental outcomes, addressing its sociodemographic classification (Feczko et al., 2021). By keeping family units intact and matching for sibling and twin pairs, the approach aimed to ensure equivalence across the samples. Subsequent analysis showed no significant differences in these variables between the samples, highlighting only minor demographic variations and identical cognitive performance.

### 2.2 Imaging acquisition and processing

The imaging for the Adolescent Brain Cognitive Development (ABCD) study was conducted on participants at 21 different locations across the United States, utilizing harmonized protocols on Siemens Prisma, Philips, and GE 3T scanners. Detailed specifics of the imaging methodology are further outlined in Casey et al. (2018). During the resting state scans, participants were instructed to keep their eyes open while viewing a passive crosshair for 20 minutes, to ensure a minimum of 8 minutes of data with low motion. These scans were performed using a gradient-echo EPI sequence, characterized by a repetition time (TR) of 800 ms, an echo time (TE) of 30 ms, a flip angle of 90°, a voxel size of 2.4 mm^3, and encompassed 60 slices. To monitor head motion, the FIRMM software was used at Siemens sites (Dosenbach et al., 2017).

Data processing adhered to the ABCD pipeline, executed by the ABCD Data Analysis and Informatics Core (Hagler et al., 2019). These procedures involved correcting T1-weighted images for gradient nonlinearity distortion and intensity inhomogeneity, followed by rigid registration to a custom atlas and segmentation via FreeSurfer to derive regions of interest (ROIs) for white matter, ventricles, and the whole brain. Resting-state images underwent a comprehensive correction process for head motion, B0 distortions, and gradient nonlinearity distortions, alongside registration to structural images using mutual information. The initial scan volumes were discarded, and voxel-wise normalization and demeaning were performed. The data were further refined by regressing out the signal from estimated motion time courses—including six motion parameters, their derivatives, squares, quadratic trends, and the mean time courses of white matter, gray matter, and whole brain plus their first derivatives. Frames exhibiting more than 0.2 mm displacement were excluded to mitigate motion contamination.

### 2.3 Measures

#### 2.3.1 Brain Measures

For the subtyping analysis, we used rs-fMRI connectivity metrics from the ABCD Data Repository, which were generated using a seed-based correlational method. Specifically, we used 247 measures derived from connectivity between 19 subcortical regions and 13 cortical networks, while 91 measures pertained to connectivity within and between the cortical networks, summing up to 338 RSFC measures. Note that the term ’none’ was used for regions not affiliated with any networks. Below is a brief description of the surface sampling, ROI averaging, and network correlation analysis. For full details on ABCD’s image processing and analysis methods, refer to (Hagler et al., 2019).

Preprocessed time courses were sampled onto each subject’s cortical surface for the surface sampling and ROI averaging analyses. Then average time courses were calculated for cortical surface-based regions of interest (ROIs) utilizing FreeSurfer’s anatomically-defined parcellations (Desikan et al., 2006; Destrieux et al., 2010), as well as a functionally-defined parcellation based on resting-state functional connectivity patterns (Gordon et al., 2016). These parcellations are resampled from the atlas-space to align with each subject’s space. Similarly, average time courses were computed for subcortical ROIs (Fischl et al., 2002). For each ROI, the variance over time was calculated, indicating the amplitude of low-frequency oscillations. For the network correlation analyses, correlation values were computed for each ROI pair, subsequently converting these into z-statistics via Fisher transformation. This approach generated summary measures of network correlation strength (Van Dijk et al., 2010). ROIs within the Gordon parcellation framework were categorized into various networks (e.g., default, frontoparietal, dorsal attention, etc.) (Gordon et al., 2016). The average correlation within a network was determined by averaging the Fisher-transformed correlations of every unique ROI pair within that network. For inter-network correlations, the correlations of every unique ROI pair between two different networks were averaged. Additionally, the correlation of each network with each subcortical gray matter ROI was assessed by averaging the correlations between every ROI in a given network and each subcortical ROI.

#### 2.3.2 Behavioral Measures

The present report focused on cognitive and emotional measures due to their potential relevance to mental health outcomes in children and adolescents. The cognitive measures, derived from the NIH Toolbox, assessed several cognitive domains highly relevant in academic and social success. Impulsivity, measured via the UPPS-P questionnaire, was chosen due to its relevance in understanding behavioral tendencies and potential susceptibility to high-risk behaviors. The Stroop measures provided a way of assessing cognitive control over emotional information. Lastly, the parent-reported psychopathology symptoms via the Child Behavior Checklist (T. M. Achenbach & Ruffle, 2000) allowed us to use more naturalistic assessments of behavioral and emotional difficulties. Together, these measures provide a robust, multi-dimensional overview of factors contributing to cognitive functioning and mental health, thus assisting in understanding the relationship between these domains and our RSFC subtypes.

##### 2.3.2.1 Cognitive and Executive Functioning

Cognitive functioning was assessed in several ways, many of which were based on tasks included in the ABCD battery that utilize portions of the National Institutes of Health (NIH) Toolbox, which assesses various cognitive domains such as memory, language, and processing speed (Bleck et al., 2013; Gershon et al., 2013; Hodes et al., 2013) . First, we used principal component scores representing three broad cognitive domains from the neurocognitive battery of the ABCD dataset (Luciana et al., 2018) that were derived using a Bayesian Probabilistic Principal Components Analysis (BPPCA) model, which accounts for site-specific and familial variations (Thompson et al., 2019). To summarize, the NIH Toolbox provided seven cognitive metrics: Picture Vocabulary evaluates language proficiency and verbal intelligence; Oral Reading Recognition assesses reading capabilities; Pattern Comparison Processing Speed gauges swift visual processing; List Sorting Working Memory evaluates memory based on categories and perceptions; Picture Sequence Memory tests memory through sequencing activities; Flanker Inhibitory Control and Attention assesses conflict processing and response inhibition; and Dimensional Change Card Sort evaluates cognitive adaptability. Beyond the NIH Toolbox, two additional tasks were considered: The Rey auditory verbal learning test (RAVLT), which assesses auditory memory and recognition, and the Little Man Task, which evaluates visual-spatial processing, especially mental rotation.

The BPPCA model identified three primary component scores: general-ability-(BPPCA), influenced mainly by oral reading, picture vocabulary, and list sorting memory tasks; executive-capability-(BPPCA), influenced by the flanker, dimensional change card sort, and pattern comparison speed tasks; and learning/memory-(BPPCA), influenced by picture sequence memory and list sorting memory tasks. For the present report, to ensure the consistency of the separate samples, the CFA and PCA scores were derived separately from Sub-Sample-1 and Sub-Sample-2 (Feczko et al., 2021).

Second, we employed a previously established EF model derived from the ABCD baseline study data (Freis et al., 2022), which is grounded in the widely supported unity/diversity model of EFs (Friedman & Miyake, 2017). This measure was derived from performance on the Flanker, Card Sort, and List Sort tasks from the NIH Toolbox2 and behavioral data from two neuroimaging tasks: the Emotional N-back and Stop-Signal tasks (SST). The reliability of these tasks has been validated and documented in pilot ABCD data and prior research (Casey et al., 2018; Luciana et al., 2018). Factor loadings for each sample are reported in **Supplemental Table 5**. The three derived factors were common-EF-(CFA), cognitive-aptitude-(CFA), and updating-specific-(CFA).

##### 2.3.2.2 Emotion-Related Processing

###### 2.3.2.2.1 Psychopathology

Parental reports were used to measure symptoms of psychopathology using the Achenbach Parent Report Child Behavior Checklist (CBCL) (T. Achenbach, 2009). The CBCL produces scores for eight scales representing different symptoms of psychopathology. These scales have shown reliability and can be used to create broader internalizing and externalizing composites (Dutra et al., 2004; Petty et al., 2008).

The current report used the eight subscale scores of the Child Behavior Checklist (CBCL) alongside the Internalizing and Externalizing psychopathology composite scales to comprehensively assess a child’s emotional and behavioral functioning. These subscales include Anxious/Depressed, which measures symptoms related to anxiety and depression; Withdrawn/Depressed, assessing social withdrawal and depressive symptoms; Somatic Complaints, focusing on physical symptoms without a clear medical cause often linked to emotional distress; Social Problems, evaluating difficulties in social interaction, including peer-related issues; Thought Problems, identifying unusual thoughts or behaviors such as strange ideas or obsessions; Attention Problems, measuring symptoms of inattention, impulsivity, and hyperactivity; Rule-Breaking Behavior, assessing behaviors that contravene accepted rules or norms, like lying, stealing, and truancy; and Aggressive Behavior, evaluating confrontational or aggressive behaviors. Additionally, the CBCL’s composite scores, “Internalizing Problems” which encompasses the Anxious/Depressed, Withdrawn/Depressed, and Somatic Complaints subscales, and “Externalizing Problems,” combining Rule-Breaking Behavior and Aggressive Behavior subscales, offer further insight into broader patterns of psychopathology. For a more in-depth description of the measures used in the ABCD study, see (Barch et al., 2021).

###### 2.3.2.2.2 Impulsivity

For the current report, we derived five dimensions of impulsivity that were measured through a child report using a condensed 20-item version of the Urgency, Premeditation (lack of), Perseverance (lack of), Sensation Seeking, Positive Urgency, Impulsive Behavior Scale (UPPS-P) (Lynam et al., 2007; Watts et al., 2020). Following the same CFA procedures outlined in Watts et al. (2020), scores for these five dimensions were derived separately from Sub-Sample-1 and Sub-Sample-2 to ensure the consistency of the separate samples and prevent data leakage across our subsequent analyses that used the saved factor scores. Factor loadings for each sample are reported in **Supplemental Table 5**.

###### 2.3.2.2.3 Emotional Word-Emotional Face Stroop

This task was designed to examine cognitive control to focus on task-relevant information when the task-irrelevant information is emotionally salient. In this task, participants categorized the emotional valence (positive, negative) of a word while disregarding an accompanying face, whose facial expression might match (congruent) or conflict with (incongruent) the word’s valence. This task, executed on an iPad, comprised two blocks: one with a 75% congruent and 25% incongruent trial split (’mostly congruent block’) and another with equal percentages of both (’equal block’). Each block consists of 48 trials, allowing 2000 ms for response. The facial stimuli from (Guyer et al., 2008) feature white adolescents expressing happiness or anger.

In the current study, we used the difference in accuracy between incongruent and congruent trials as our measure of performance, calculated separately for trials in which the distracting face was happy and those in which it was angry. This measure indicates the participant’s ability to manage cognitive interference and maintain task focus. For further details on this task and its implementation, refer to Smolker et al. (Smolker et al., 2022).

### 2.4 Procedure

#### IVEPR: Identification and Validation

##### 2.4.1 Precise Subtyping with Bagging-Enhanced Leiden Community Detection

The foundation of our analysis was established through bagging-enhanced Leiden Community Detection (LCD) (DeRosa, Kim, et al., 2023; Traag et al., 2019), a data-driven clustering methodology. LCD is an effective strategy for identifying distinct subgroups, in the current case, with analogous functional connectivity properties. Each sample was individually processed using this procedure, which was applied to both cortical and subcortical resting-state functional connectivity (RSFC) measures to identify the different subtypes. The strength of the bagging-enhanced LCD lies in its ability to address uncertainty inherent in the input data, in this case, the connectivity matrix, through bootstrapping. Additionally, its data-driven nature lends itself well to exploratory analyses, bypassing potential biases from preconceived assumptions about the number of subtypes or their structure (Nikolaidis et al., 2022, 2021). The validation stage confirms the discovered subtypes’ reproducibility, reliability, and robustness. We employed a split-sample approach, which used comparisons between subtypes and their profiles in divided samples for initial validation. This comparison is a stepping-stone for further confirmation of subtype robustness and reliability.

The LCD algorithm is a data-driven method that identifies optimal communities within a complex network, such as the network of brain regions we examined. At its core, the LCD algorithm optimizes a metric called modularity (Q), a measure of the strength of the division of a network into communities. It compares the density of connections within communities versus those between them. High modularity implies many connections within communities and only a few between them, which means the divisions are well-defined. The algorithm begins with every node (in our case, individuals) in its own community. Then, it iteratively evaluates the impact on modularity by moving each node to a different or merging community. The move or merge that maximizes the modularity is performed, and the process is repeated until no further improvements can be made, reaching a locally optimal community structure. This structure, wherein no single move or merge can enhance modularity, represents the optimal number of communities (clusters) for that run. However, how nodes are evaluated for potential moves or merges can influence the LCD algorithm’s solution. Refer to DeRosa et al. (2023) for additional information regarding this procedure.

#### IVEPR: Validation

##### 2.4.2 Employing Gradient Boosted Decision Trees and Shapley Additive Explanations for Neurodevelopmental Subtype Classification and Feature Importance

We conducted an out-of-sample classification accuracy assessment to evaluate the reproducibility of our identified subtypes using Sub-Sample-1 as the training set and Sub-Sample-2 as the testing set. Our classification process used the gradient-boosted decision trees method via the XGBoost algorithm (Chen & Guestrin, 2016). A gradient-boosted decision tree is a machine-learning technique that optimizes prediction accuracy by combining multiple weak decision trees through iterative improvement and error correction. This XGBoost algorithm is notable for its efficacy in handling high-dimensional data sets, such as neuroimaging data.

A distributed hyperparameter optimization process was carried out to achieve optimal model performance using the Ray Tune library in Python (Liaw et al., 2018). This package efficiently searches the hyperparameter space using a cross-validation-based approach and returns the optimal parameters that minimize the loss function. The optimized hyperparameters were then used to fit the final XGBoost classifier on the training data. The XGBoost algorithm was implemented with the objective function “multi:softmax”, indicating a multi-class classification problem. The tree method was set to “hist”, which uses histogram-based algorithms to grow trees. Finally, the XGBoost classifier was applied to the holdout data to generate predictions.

To further understand the model classifications, we used Shapley additive explanations (SHAP) to quantify the contribution of each feature to the prediction (Lundberg & Lee, 2017). SHAP values were computed for the holdout data using a tree explainer. To reveal the most important features driving the classification, the model’s feature importances were extracted and ranked. The top 15 features were selected and used to visualize the subtypes. This analysis provided a comprehensive understanding of the identified subtypes’ classification based on the RSFC features and offered insights into the most influential features in the classification process.

Finally, to validate the authenticity of these subtypes, we took measures to confirm they were not solely influenced by socioeconomic status (SES) and demographic factors which were household income, parental marital status, parental education, marital status, and adversity. To do this, we used the SES and demographic features as predictors for the subtypes. This step was necessary to ensure the identified subtypes were not predominantly the result of underlying socio-economic or demographic indicators.

#### IVEPR: Evaluation

##### 2.4.3 Assessment of Cognitive and Executive Functioning and Mental Health Using Latent Models and Confirmatory Factor Analysis

We performed confirmatory factor analysis (CFA) to derive two latent model factors representing a) cognitive and executive functioning and b) impulsivity. We used a task-based latent variable model of Executive Functions (EFs) from Freis et al. (2022) for the first model. For the second, we used the self-reported Urgency, Premeditation (lack of), Perseverance (lack of), Sensation Seeking, and Positive Urgency subscales from the Impulsive Behavior Scale (UPPS-P) as outlined by Barch et al. (2018). Age and sex were regressed out of all measures before extracting the latent factors and using these scores for subsequent analyses. Furthermore, we conducted invariance testing to validate the construct measurement and ascertain that the detected differences across subtypes genuinely represented underlying differences in the constructs of interest (Millsap, 2011). Verifying measurement invariance across our constructs ensures the consistent representation of the same constructs across varied subtypes. All analyses, including latent model derivation and CFA invariance testing, were conducted using the Lavaan package in R (Rosseel, 2012).

##### 2.4.4 Neurodevelopmental Differences in Cognitive and Executive Functioning and Mental Health

To evaluate differences in cognitive and executive functioning, impulsivity, and psychopathology across the identified subtypes, we used ANOVA with false discovery rate (FDR) correction (Benjamini & Hochberg, 1995). Both split samples (Sub-Sample-1 and Sub-Sample-2) and the Full-Sample, which combined the subtypes from Sub-Sample-1 and Sub-Sample-2, were subjected to this analysis. ANOVA was conducted to determine if there were any statistically significant differences among the means of the subtypes for cognitive and executive functioning, impulsivity, and psychopathology. Following the ANOVA, FDR-corrected post-hoc comparisons were performed to identify which specific subtypes were significantly different from each other concerning cognitive and executive functioning, impulsivity, and psychopathology. We first identified significant subtype group-level differences across both samples. We then deemed differences to be reproducible if at least one pairwise post hoc comparison was significant in both Sub-Sample-1 and Sub-Sample-2 for cognitive and executive functioning, impulsivity, and psychopathology measures. This criterion ensured that the observed differences were consistent across both split samples.

#### IVEPR: Prediction

##### 2.4.5 Evaluating Subtype Importance in Brain-Behavior Predictive Models

Our Brain-Behavior predictive modeling pipeline began with a systematic approach to RSFC before computing the conditional random forest (CRF) models that included the categorical Subtype feature as a predictor. A conditional random forest is an advanced machine learning model that builds multiple decision trees to make predictions, adjusting for specific conditions or variables to capture complex interactions and dependencies within the data. We implemented the BorutaShap method (Kursa & Rudnicki, 2010) using the Boruta-Shap package in Python to perform this feature selection. Unlike traditional approaches that lean heavily on the inherent feature importance of random forests, BorutaShap capitalizes on SHAP values to determine feature importance, establishing it as a more model-agnostic method. For each of our cognitive functioning and mental health measures, our process involved fitting an XGBoost model, followed by the computation of SHAP values for each feature. The Boruta algorithm was subsequently applied. This approach is particularly advantageous for pre-processing steps in CRF models. Two primary reasons underpin this assertion: 1) Dimensionality Reduction - which optimizes training times by reducing irrelevant features and omitting noisy features; and 2) Compatibility - Boruta’s algorithm was initially designed for random forests, making it congruent with models like CRFs.

Building on this, we sought to discern whether the RSFC subtypes consistently outperform the RSFC features as a predictor for cognitive and mental health scores using CRFs from the party package in R (Strobl et al., 2008). There are several reasons why CRFs are apt for gauging feature importance and assessing the predictive capacity of neuroimaging-based subtypes. Importantly, CRFs can accommodate categorical features with multiple levels, such as our Subtypes, and are favored over XGBoost, which necessitates transforming categorical measures into dummy-coded features for model inclusion. Unlike parametric models, CRFs are not tethered to assumptions about data distributions, enabling them to delineate intricate, non-linear relationships between predictors and outcomes. Such adaptability is crucial when analyzing high-dimensional neuroimaging data. CRFs also leverage the inherent structure of decision trees to assess multivariate patterns across the included measures, allowing for a comprehensive evaluation of how interactions among variables contribute to predicting cognitive and mental health outcomes. Additionally, the ensemble nature of CRFs offers resistance to overfitting. CRFs also harness a “bagging” strategy, wherein multiple trees constructed from random data subsets are aggregated, promoting model stability and generalizability. Finally, CRFs allow for an interpretable measure of feature importance through “permutation importance”, which quantifies the dip in prediction accuracy when a feature is randomly shuffled.

To bolster the robustness of our findings, we performed 1000 iterations of the CRFs for each cognitive functioning and mental health measure. Each measure’s feature importance was ranked across these iterations. Subsequently, for both Sub-Sample-1 and Sub-Sample-2, we calculated the mean feature importance ranks across the thousand iterations. Keeping the samples separate was essential to gauge the consistency of top features in each sample, preventing undue influence of one sample’s features on the other. The resultant feature importance means from both samples were averaged to procure a definitive ranking by measure.

To evaluate the significance of the Subtype feature across our CRF models, we focused on its ranking in terms of feature importance across all the measures. Specifically, we calculated how frequently the Subtype feature appeared as the most important predictor in different ranking tiers: top 1, top 5, and top 10. This analysis involved calculating the proportion of times the Subtype feature was ranked as the most important (top 1), within the five most important (top 1-5), and the ten most important features (top 1-10) for predicting each measure. We then conducted similar proportion calculations for each of the individual RSFC features, assessing their rankings in the top 1, top 5, and top 10 positions. This approach allowed us to directly compare the subtype’s predictive ability against other RSFC features. Our final assessment determined whether the Subtype consistently emerged as the leading predictor across these three tiers of feature importance. By juxtaposing the Subtype’s rankings against the RSFC features, we sought to ascertain its relative importance in forecasting various cognitive and mental health outcomes.

#### IVEPR: Replication

##### 2.4.6 Assessing Reproducibility, Differences, and Predictive Ability of Neurodevelopmental Subtypes

We used two key strategies, bootstrapping and split-sample subsampling, for the robust assessment of the reproducibility of the identified subtypes, their differences, and their predictive ability. For additional details on the rationale of our replication analyses, refer to the **Supplemental Materials.**

###### Bootstrapping

The first strategy was bootstrapping. Sub-Sample-1 and Sub-Sample-2 were first partitioned to maintain an equal number of individuals within each subtype. This procedure was implemented to protect against increasing the imbalance in individuals from each subtype drawn during the bootstrap resampling. Each partitioned sample was then resampled with replacement (bootstrapping), with each bootstrapped iteration comprising 66% of the individuals from the original sample. We performed ANOVA on each new bootstrapped sample’s cognitive functioning and mental health measures. This process was repeated for 1000 iterations for each bootstrapped sub-sample. Furthermore, we resampled the Full-Sample of individuals who passed RSFC quality control with replacement and ran it through the ANOVA analyses. The robustness and reliability of a given measure were assessed by extracting the FDR-corrected p-values from each iteration and ensuring their mean value was less than .05 across all three samples (Sub-Sample-1, Sub-Sample-2, Full-Sample).

###### Split sample down-sampling

The second strategy we used was split-sample down-sampling. For each of the nine down-sampling increments (from 10% to 90% of the total), we generated 100 new samples, resulting in a total of 900 samples (9 increments × 100 samples). Since we had 200 new samples for Sub-Sample-1 and Sub-Sample-2, this amounted to 1800 new samples (900 samples per split × 2 splits = 1800 samples). Bagging-enhanced LCD was applied to these 1800 new samples to derive the down-sampled subtypes. We then performed a series of tests on each of the 1800 new down-sampled subtypes, including calculating modularity (Q), the Adjusted Rand Index (ARI) compared to the complete set of individuals within a given subtype for each of Sub-Sample 1 and Sub-Sample 2, the mean maximum correlations to full split-samples, and the maximum correlation across each of the other subtypes within the 100 iterations. Specifically, the mean maximum correlations were calculated by comparing each down-sampled subtype’s connectivity patterns to the corresponding original complete set of individuals for a given subtype within each of the two sub-samples. The ARI (Adjusted Rand Index) is a statistical tool used to evaluate the similarity between two sets of subtypes, factoring in the likelihood of random chance. It effectively quantifies the consistency of the identified subtypes across different sub-samples, providing a solid foundation for evaluating the quality and reproducibility of the subtype identification process.

Additionally, these down-sampled subtype profiles were compared across the two sub-samples, assessing how these down-sampled subtypes reliably replicate the connectivity profiles observed in the larger, complete original sub-samples and maintain consistency across different subsets. We evaluated the success, reproducibility, and reliability based on the mean maximum correlations between the original subtype profiles in each split sub-sample and the down-sampled subtype profiles. Furthermore, we calculated the ARI to the original sample subtype label. We defined success as high mean maximum correlations (average > .9) across the subtypes for each down-sampling split sample, as well as to the original labels. We also expected the Adjusted Rand Indexes to outperform chance for each sample comparison and to show an increasing trend as the sample size grows.

This sub-sampling approach aims to rigorously assess the reproducibility and reliability of subtype detection across varying sample sizes, directly addressing potential concerns when dealing with smaller samples in future studies. Importantly, achieving high reproducibility in these down-sampled datasets is a prerequisite for advancing to the second objective of our methodology. Only upon validating the stability and consistency of our subtype models through this down-sampling approach could we confidently apply these models to new samples without re-clustering. Such a step is crucial for demonstrating the generalizability of our findings. This conditional progression underlines the importance of our initial down-sampling strategy in ensuring that our models possess the robustness required for effective application to diverse populations. Success in this first phase would enhance the translational potential of these RSFC subtypes, mirroring the approach used in developing polygenic scores from large discovery samples and emphasizing the utility of large discovery sets for accurately applying neuroscientific models across varied demographic contexts.

##### 2.4.7. Evaluating the Robustness of RSFC Subtypes Against Noisy Data

We had two primary reasons for these supplementary subtyping analyses. First, we aimed to determine if including individuals with noisy fMRI data would impact the subtypes derived from those with cleaner data. To achieve this, we executed the same LCD analyses on the complete sample (N=9027) of ABCD participants with neuroimaging data, ensuring only one sibling was selected from each family. Subsequently, we calculated the maximum correlations across the subtypes for both samples. While one might object to including individuals with noisy data, there is evidence that certain cognitive variables, such as executive function, may correlate with head motion (Wylie et al., 2014). If that is the case in this sample, excluding them might influence the brain-behavior relationships between the identified subtypes. Hence, we wanted to test whether our results are robust against such potential bias. However, this question’s validity hinged on whether the subtype profiles were consistent across the entire and include-only samples. To assess this, we performed bootstrapped ANOVAs to gauge the consistency of subtype differences across both groups (see **Supplemental Table 6** and **Supplemental Figure 3**).

Furthermore, we aimed to ascertain if the subtypes would be consistent in the “high motion” sample (N=1,293). It could have significant implications if these subtypes are reproducible within that group. It might allow us to retain these individuals in future analyses, reinforcing a primary objective of our paper: to demonstrate that the subtype (i.e., whole-brain profile) might be a more reliable and meaningful neuro-marker than individual RSFC connections. To test this idea, we employed the same LCD analyses to evaluate the consistency of subtype profiles. The outcomes of these supplementary analyses are briefly discussed in the results and discussion sections, with more detailed results available in the supplemental materials section.

### 2.5. Code Accessibility

Custom Python, R, and bash code for all primary statistical analyses are available at https://github.com/jakederosa123/neuro_dev_rsfc_subtypes_abcd-

## 3. Results

### IVEPR: Identification and Validation

This study employed bagging enhanced LCD on RSFC data to identify distinct neurodevelopmental subtypes and examine their relationships with demographic and behavioral traits. Specifically, four subtypes were identified via LCD on the RSFC data. All subtypes showcased high reproducibility (r range=0.98-0.996) and strong associations with demographic and behavioral measures. We conducted an out-of-sample classification accuracy assessment to assess the reproducibility of our identified subtypes across Sub-Sample-1 and Sub-Sample-2. This analysis yielded an accuracy of 88.90%, underscoring reasonable robustness and reproducibility of the subtypes across the two samples. To ensure that our subtypes were not merely artifacts influenced by SES and demographic factors, we used these SES and demographic features as predictors for the subtypes. This analysis yielded a low accuracy of 32.35%, suggesting that SES and demographic factors do not predominantly drive these subtypes.

Regarding the distribution of individuals across the identified RSFC subtypes, we found a uniform presence of all four subtypes across the different samples, affirming their broad applicability within the population (**Supplemental Table 1**). Specifically, Subtype-1 emerges as the most prevalent, with its presence marked by 26.81% in Sub-Sample-1, 28.35% in Sub-Sample-2, and 28.27% in the Full-Sample. Subtype-2 also exhibits a notable representation, encompassing approximately a quarter of each sample (25.29% in Sub-Sample-1, 22.44% in Sub-Sample-2, and 24.35% in the Full-Sample), reinforcing the diversity and consistency of neurodevelopmental patterns in the population. Subtypes 3 and 4, while showing slightly lower percentages, particularly in Sub-Sample-1 (21.93% for Subtype-3 and 25.97% for Subtype-4) and the Full-Sample (22.56% for Subtype-3 and 24.82% for Subtype-4), nonetheless maintain a significant presence. This distribution aligns with the assumed nested heterogeneity within the ABCD sample, where each subtype, characterized by unique RSFC patterns, coexists within the population.

The demographic and phenotypic associations reported below were replicated across the two independent samples (Sub-Sample-1 and Sub-Sample-2), meaning that significant FDR-corrected differences that involved the same variables and directions were observed across both samples. For each subtype, we begin by characterizing the prominent features of the respective imaging profile, followed by the prominent features characterizing their demographic and phenotypic profiles. Refer to **Supplemental Table 2** for a comprehensive report on all phenotypic and **Supplemental Table 1** demographic comparisons across the subtypes, **Figure 3** for the RSFC profiles of each subtype, and **Figure 4A-B** for the cognitive and mental health profiles for each subtype by sub-sample.

**Figure 3.**
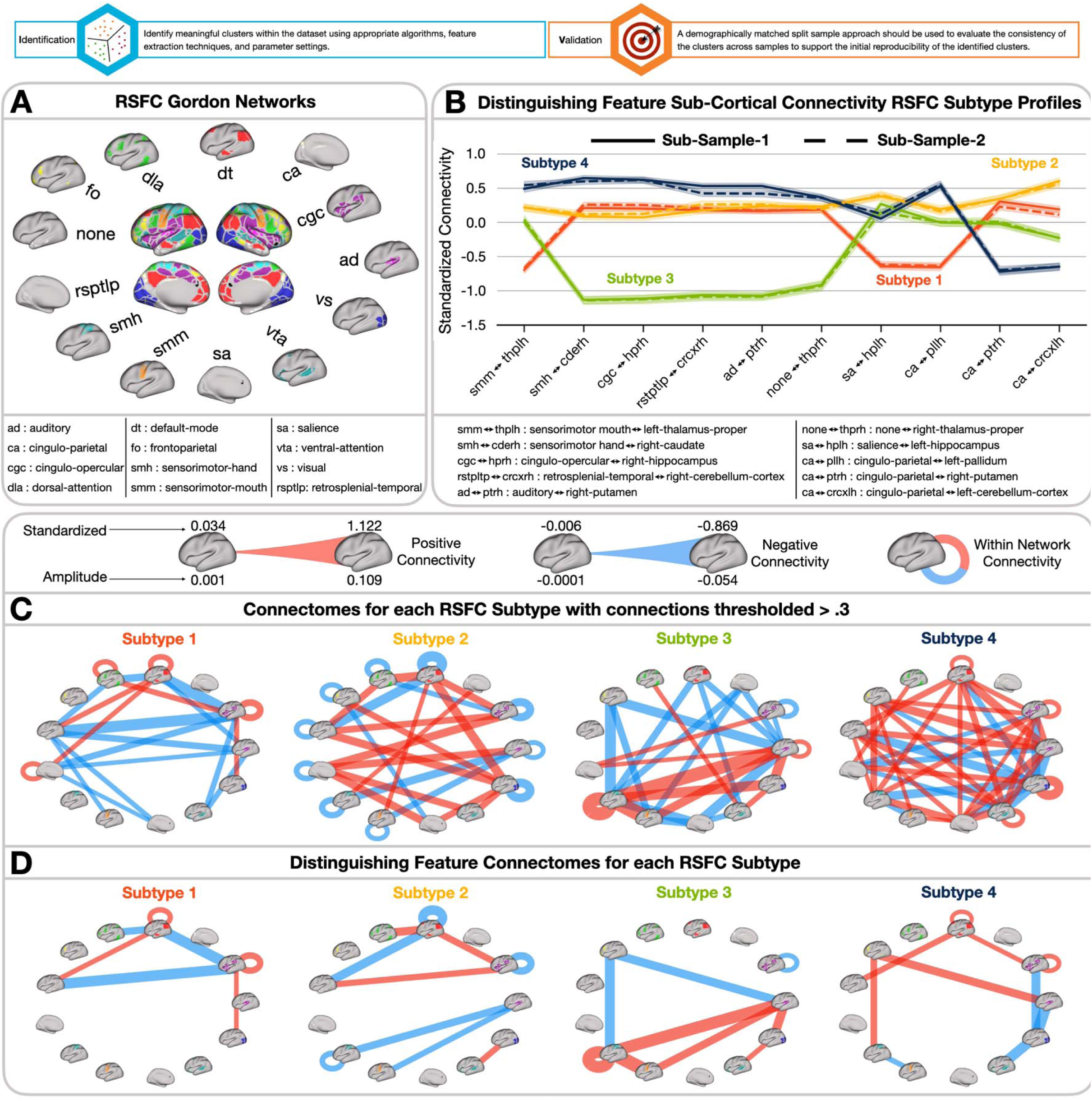
Resting State Functional Connectivity (RSFC) Subtype Profiles. **A)** RSFC Gordon Networks legend for **C** and **D.** All 13 networks are displayed and labeled accordingly. **B)** Subtype profiles represent the mean standardized functional connectivity among the top 10 sub-cortical regions identified by SHAP feature importances for classifying each subtype. Lines are colored by Subtype association (Subtype-1, orange; Subtype-2, yellow; Subtype-3, green; Subtype-4, blue) and differentiated by sample (Sub-Sample-1, straight, Sub-Sample-2; dashed). Full sub-cortical ROI and cortical network names are displayed below the profiles. For both **C** and **D,** the line thickness represents the connectivity strength. Connectivity directionality is denoted by blue for negative and red for positive. Self-loops characterize within-network connectivity. Given the high reproducibility across Sub-Sample-1 and Sub-Sample-2 subtypes, the Full-Sample subtypes are displayed from left to right. **C)** Subtype profiles represent the functional connectivity among cortical regions based on a connectivity threshold of 0.3 for each RSFC Subtype. **D)** Subtype profiles represent the functional connectivity among the top 15 cortical regions identified by SHAP feature importances for classifying each RSFC Subtype. Note: The amplitude of connections by Subtype can be seen in **Supplemental** Figure 1.

**Figure 4.**
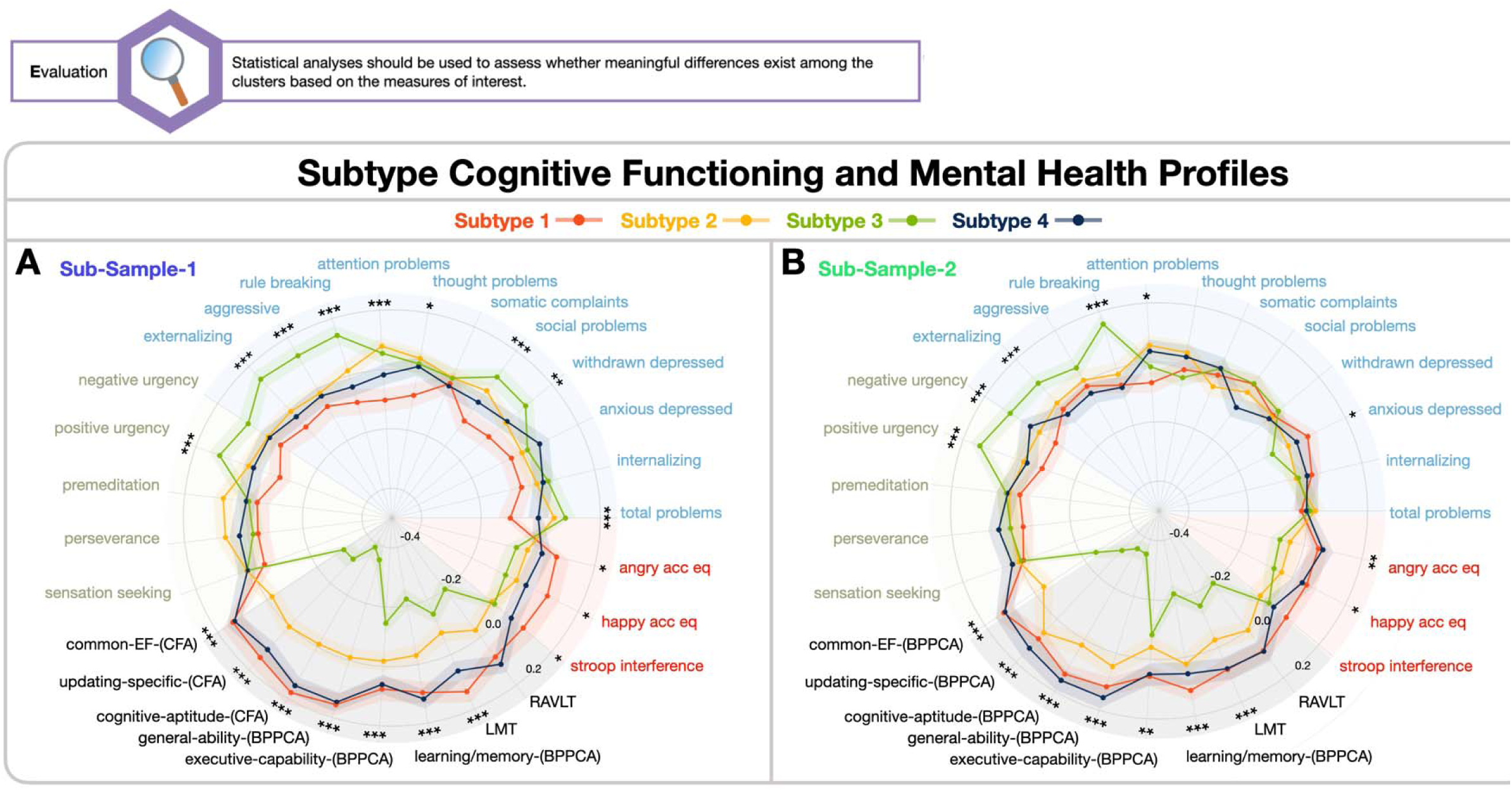
RSFC Subtype Cognitive and Mental Health Profiles. Illustration of the mean residualized z-score for different phenotypic measures, colored by subtype (Subtype-1, red; Subtype-2; yellow; Subtype-3, green; Subtype-4, blue) by Sub-Sample-1 **A)** and Sub-Sample-2 **B)**. Points represent the mean age and sex residualized z-score for each cognitive functioning and mental health measure. Shaded bars signify each point’s 95% confidence interval and are colored by Subtype. Sectors of the radar plots are color-shaded by domain allegiance (impulsivity; green; psychopathology, blue; emotion and cognition; red; cognitive functioning; black). Refer to Figure 3 B-D for Subtype patterns of connectivity.

In comparisons between the “passed RSFC quality control” sample and the “complete sample”, the subtype maximum correlations consistently exceeded .99. In comparisons between the “passed RSFC quality control”, “complete sample” sample, and the “high motion” sample, the subtype correlations varied between .623 and .965. It is worth highlighting that subtypes 1 and 3 in the “ high motion” sample exhibited the highest correlations to the “passed RSFC quality control” and “complete sample” subtypes 1 and 3, both surpassing .9. See **Supplemental Figure 2** to view these correlations and RSFC profiles across the different inclusion criteria sample subtypes and **Supplemental Tables 3-4** for subtype demographics.

Finally, we conducted additional analyses to verify that our identified subtypes were not artifacts of frame displacement (FD) (i.e., head motion). In these analyses, we removed the influence of FD from the RSFC data before generating the subtypes. The resulting subtype profiles demonstrated high reproducibility between Sub-Samples 1 and 2, with maximum correlation values ranging from .96 to 1. Furthermore, most individuals remained in their respective subtypes, as indicated by the adjusted rand indices (ARI) of .78 for Sub-Sample-1 and .85 for Sub-Sample-2. These findings suggest that FD did not substantially affect the subtypes we initially identified.

### IVEPR: Evaluation

#### 3.2.1. RSFC Subtype Profiles

##### Subtype-1

Subtype-1 is characterized by strong positive within default mode network connectivity and strong negative connectivity between auditory and sensorimotor-hand networks and between the cingulo-opercular and default mode network. This subtype also shows strong negative connectivity between the default mode and the dorsal attention network. Furthermore, this subtype exhibits strong positive connectivity between the cingulo-opercular network and the left caudate and right hippocampus while displaying strong negative connectivity with the right ventral diencephalon. In addition, the sensorimotor-hand networks demonstrate strong negative connectivity with the sensorimotor-mouth networks, strong positive connectivity with the left pallidum and right caudate, and strong negative connectivity with the right hippocampus.

Subtype-1 performed better on all cognitive and EF than Subtypes 2 and 3. Subtype-1 also did not reveal a high degree of mental health problems. Regarding demographics, children in Subtype-1 predominantly come from families where parents have a high level of education, with many holding post-graduate degrees. Most families in this subtype have upper-range household incomes, and a majority of their parents are married. Subtype-1 was also characterized by relatively lower adversity scores than the other subtypes. The gender distribution is nearly equal between males and females.

##### Subtype-2

Subtype-2 is characterized by strong negative connectivity between auditory and sensorimotor-hand networks and strong positive connectivity within the cingulo-opercular network and with the default mode network. Additionally, individuals in this group show strong positive connectivity between the cingulo-opercular network and the left caudate and right hippocampus, along with strong negative connectivity between the sensorimotor-hand networks and sensorimotor-mouth networks. This subtype also shows robust negative within-network connectivity in all networks except the salience network.

Compared to other subtypes, Subtype-2 performed poorly across various cognitive measures, including the LMT, RAVLT, general capability, executive-capability-(BPPCA), and learning/memory-(BPPCA). For the Cognitive and EF latent factors, Subtype-2 performed lower than Subtype-4 but performed better than Subtype-3 in common-EF-(CFA), cognitive-aptitude-(CFA), and the Updating-Specific factor. Demographically, children in Subtype-2 come from families whose parents have varied educational backgrounds, though a significant proportion have bachelor’s and post-graduate degrees. Household incomes in this subtype are varied but tend to be relatively high, and most parents are married. Subtype-2 children had significantly higher adversity scores than Subtype-4 but lower than Subtype-3. There is an even gender distribution in Subtype 2.

##### Subtype-3

Children in Subtype-3 exhibit strong positive connectivity between auditory and sensorimotor-hand networks and moderate positive connectivity within the cingulo-opercular network. They also display pronounced negative connectivity between the frontoparietal and sensorimotor-hand networks and the frontoparietal and auditory networks. Notably, there are strong negative connections between the cingulo-opercular network and the left caudate, right hippocampus, and right ventral diencephalon for this subtype. Furthermore, the sensorimotor-hand networks demonstrate strong positive connectivity with sensorimotor-mouth networks while displaying negative connections with the left pallidum, right caudate, and right hippocampus.

Notably, these children performed worse in most all cognitive and EF measures compared to the other subtypes. Subtype-3 also had higher externalizing and rule-breaking behavior problems on average. Demographically, the parents of children in Subtype-3 tend to have varied educational backgrounds, with a significant portion having some college education. This subtype includes many families with lower household incomes and fewer parents who are married compared to the other subtypes, and they have higher adversity scores. The gender distribution is approximately even between males and females.

##### Subtype-4

The children in Subtype-4 exhibit weak negative connectivity between auditory and sensorimotor-hand networks and negative connectivity between the default mode and dorsal attention network, similar to Subtype-1. They also show positive connectivity within the default mode and cingulo-opercular connectivity with the left caudate, right hippocampus, and right ventral diencephalon. Additionally, the sensorimotor-hand networks display negative connectivity with sensorimotor-mouth networks while demonstrating positive connectivity with the left pallidum, right caudate, and right hippocampus.

Similar to Subtype-1. Subtype-4 outperformed Subtype-2 and Subtype-3 on all cognitive and EF measures. Subtype-4 also did not reveal major mental health problems. Demographically, a significant number of parents of the Subtype-4 children hold post-graduate degrees. The families of children in this subtype generally have higher household incomes, and many of the parents are married. Subtype-4 revealed lower adversity than Subtypes 2 and 3 and equivalent to Subtype-1. There is a nearly even distribution of genders, with a slightly higher number of males.

#### 3.2.6. Invariance Testing across Factors Underlying Cognitive and Mental Health Difficulties

We evaluated metric invariance across the RSFC subtypes using fit indices and Chi-square difference tests for the two models (EF model and the UPPS-P factor model). We observed evidence supporting the establishment of minimum metric invariance in all three-factor models, as indicated by acceptable fit indices and the absence of significant Chi-square differences upon adding constraints, which were consistently observed across all models. Refer to **Supplemental Table 5** for a complete report of invariance testing results.

### 3.3. Evaluating Subtype Importance in Brain-Behavior Predictive Models

To evaluate the significance of our Subtypes in predicting the 27 cognitive functioning and mental health measures compared to the individual RSFC connections, we performed 27 CRF models for each sample. These measures were selected to cover a broad spectrum of cognitive abilities and mental health-related conditions. Analyzing the feature importance ranks from these models across both samples, we found that, out of the 27 measures, the Subtype emerged as the top predictor for 5 measures (18.52%). It also ranked among the top 5 predictors for 11 measures (40.74%) and the top 10 for 17 measures (62.96%). Notably, the Subtype consistently secured its position within the top 1, 5, and 10 features across Sub-Sample-1 and Sub-Sample-2 (see **Figure 5**). For a detailed breakdown of the counts and proportions of the most influential RSFC connections that ranked among the top 1, 5, and 10 predictors, see **Supplemental Table 6**.

**Figure 5.**
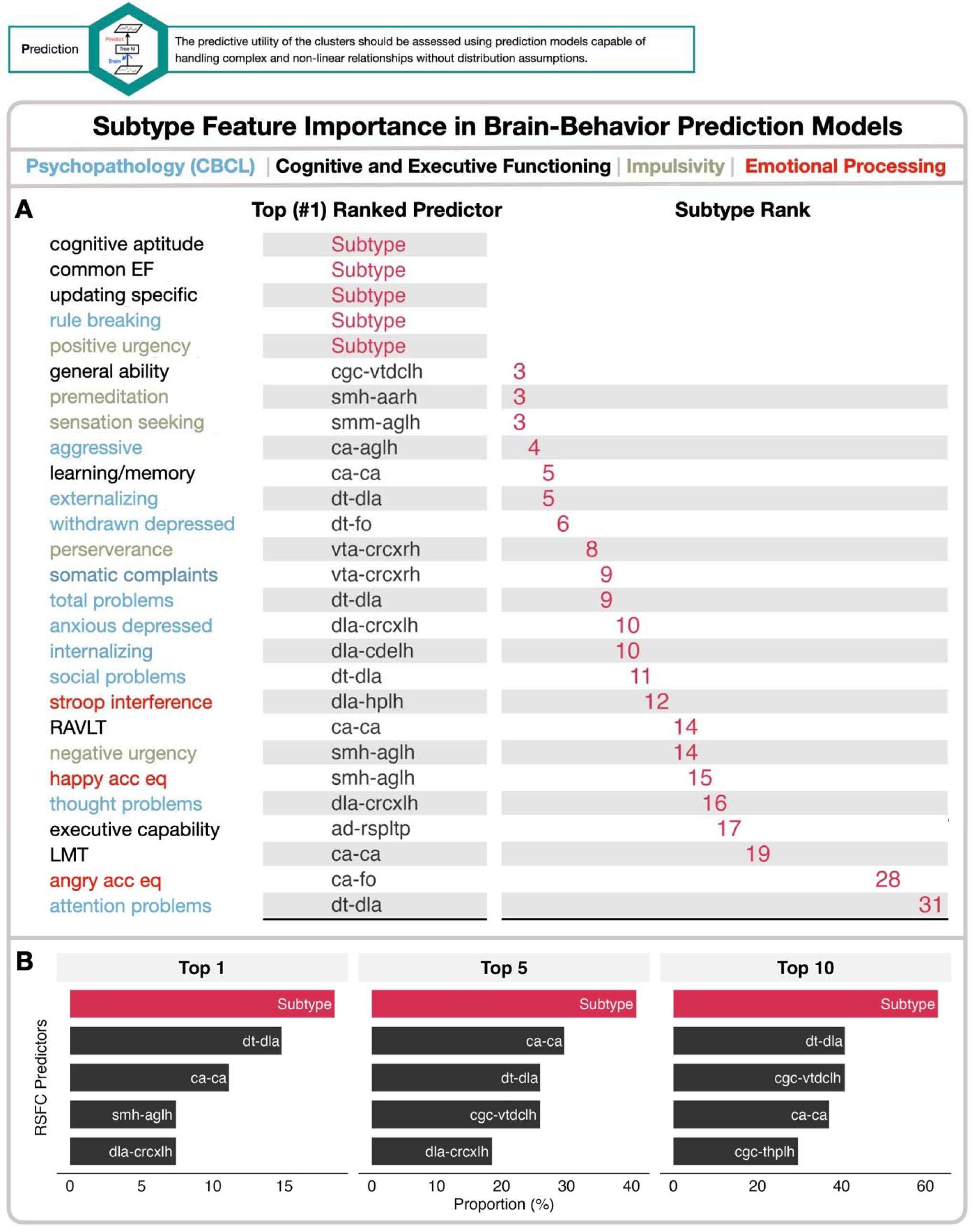
Subtype feature importance in brain-behavior prediction models. **A)** Top predictor and Subtype rank across all 27 cognitive functioning and mental health brain-behavior prediction models. **B)** The proportion of times the RSFC Subtype and connections ranked in the top 1, 5, and 10 features out of the 27 brain behavior prediction models. Note: The subcortical ROIs abbreviations and respective names are: crcxlh: left cerebellum cortex, aglh: left amygdala, crcxrh: right cerebellum cortex, vtdclh: left ventral diencephalon, cdelh: left caudate, hplh: left hippocampus, aarh: right accumbens area, thplh: left thalamus proper, vtdcrh: right ventral diencephalon, pllh: left pallidum. The three derived factors from the analysis correspond to distinct models and conceptual frameworks: common EF, cognitive aptitude, and updating specific factors are associated with the CFA models; general capability, executive capability, and learning/memory components are linked to the BPPCA model.

### 3.4. How reproducible, reliable, and robust are these RSFC subtypes?

#### IVEPR: Replication

##### 3.4.1 Bootstrapped ANOVAs

Our bootstrapped ANOVA analyses, conducted on 27 measures related to cognitive functioning and mental health, revealed consistent patterns that underscore the reliability of differences among subtypes across these variables.

Refer to **Figure 6** and **Supplemental Table 7** for complete reporting of the bootstrapped outputs. Most measures that showed statistically significant differences in the original non-bootstrapped ANOVAs remained significant in the Full-Sample of individuals who passed RSFC quality control and one of the sub-samples in the bootstrapped ANOVA analyses. On average, the FDR corrected p-value across the iterations was greater than .05. Of note, the most robust effects, which were those that were significant across both sub-samples and the Full-Sample, were observed for all cognitive measures, except for executive-capability-(BPPCA), as well as positive urgency and rule breaking (see pink bars in Figure 6), Compared to the measures in the mental health domains, the cognitive functioning measures exhibited less variability in the effect size differences across the subtypes, indicating that these measures are particularly insensitive to the characteristics of the sample. In contrast, only

**Figure 6.**
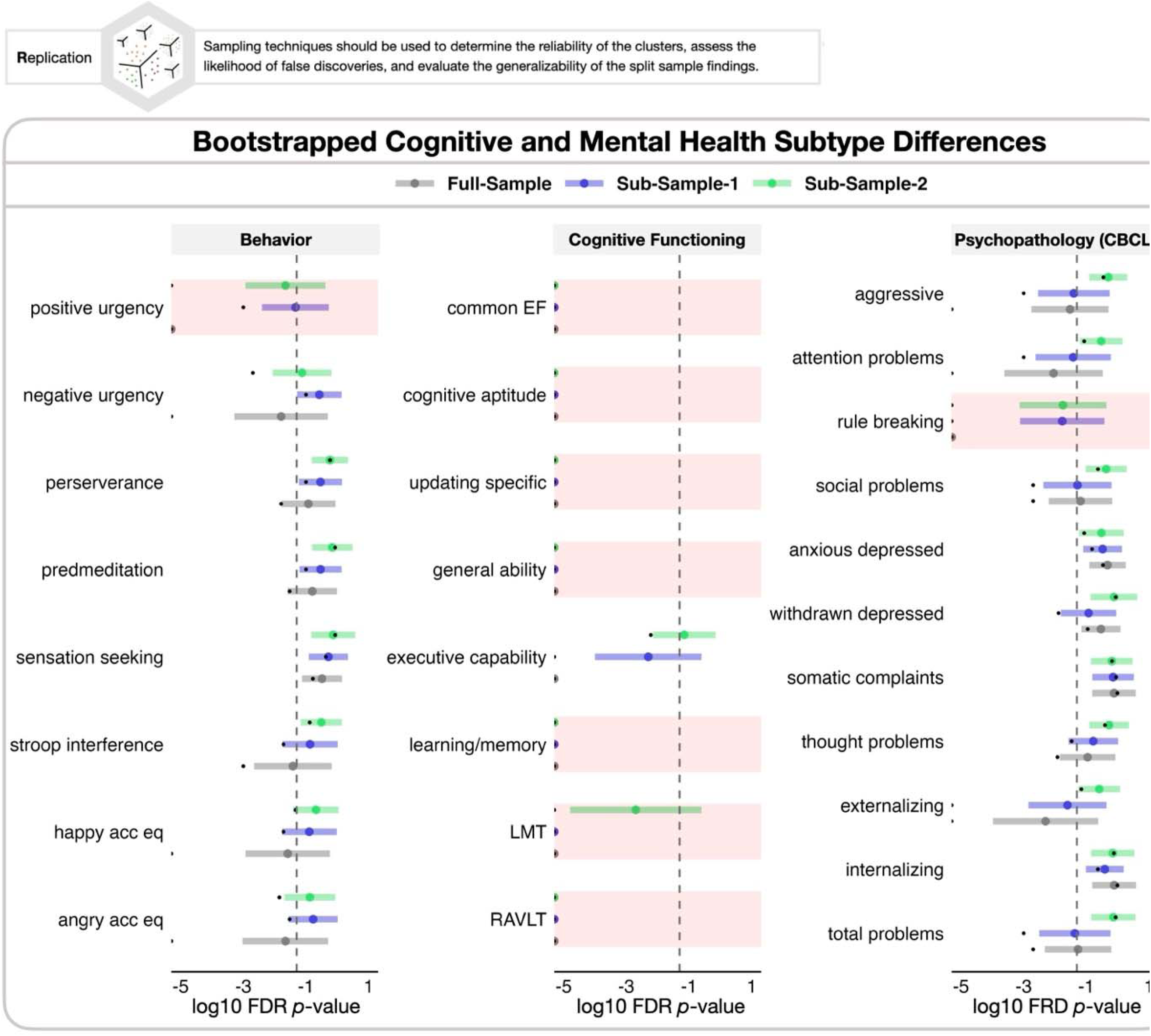
Bootstrapped ANOVAs by Sample by Phenotypic Measure. Bootstrapped ANOVAs, log10 FDR corrected P-values from each iteration are evaluated by sample, where black dots indicate the original FDR corrected p-values from the non-bootstrapped samples, colored dots and CI’s are colored by sample (Full-Sample, grey; Sub-Sample-1, blue; Sub-Sample-2; green) and represent the mean and range of FDR corrected p-values for each sample across the 1000 bootstrapped ANOVAs, and shaded bars in red indicate the mean of all three samples are < .05. Note: The three derived factors from the analysis correspond to distinct models and conceptual frameworks: common EF, cognitive aptitude, and updating specific factors are associated with the CFA models; general capability, executive capability, and learning/memory components are linked to the BPPCA model.

Regarding the bootstrapped results derived from the complete sample, we observed even more consistent reproducibility patterns than the “passed RSFC quality control” sample result reported above. Notably, the Stroop happy and angry accuracy, executive-capability-(BPPCA), and attention problems emerged as significant across both Sub-Sample-1 and Sub-Sample-2 across these analyses. See **Supplemental Figure 3** for these results.

##### 3.4.2 Split-Sample Reproducibility

As another way of evaluating our subtype’s reproducibility and reliability, we created restricted down-samples in 10% increments of each of Sub-Sample-1and Sub-Sample-2 and considered the mean maximum correlations between the original subtype profiles in each split sub-sample (Sub-Sample-1, Sub-Sample 2) and their restricted sample. We defined success based on high mean maximum correlations (average > .9) across the subtypes for each down-sampled split sample based on the original subtype labels. Our results showed consistent success as most of our down-sampled categories exhibited mean maximum correlations exceeding 0.9, especially in the down-samples that represented a higher percentage of the total sample. The ARIs demonstrated reasonable and consistent results across both samples, reinforcing the robustness of our subtypes. Importantly, even at smaller percentages like the 10% sample size, the relatively high ARI values underscore the reproducibility of the subtypes. The fact that we see consistent and reasonable ARI values at such small sample sizes and a steady increase thereafter offers compelling evidence that the subtypes are robust and can be reliably replicated across varying sample sizes. See **Figure 7** for a visual representation of these results, and a detailed breakdown is available in **Supplemental Table 8**.

**Figure 7.**
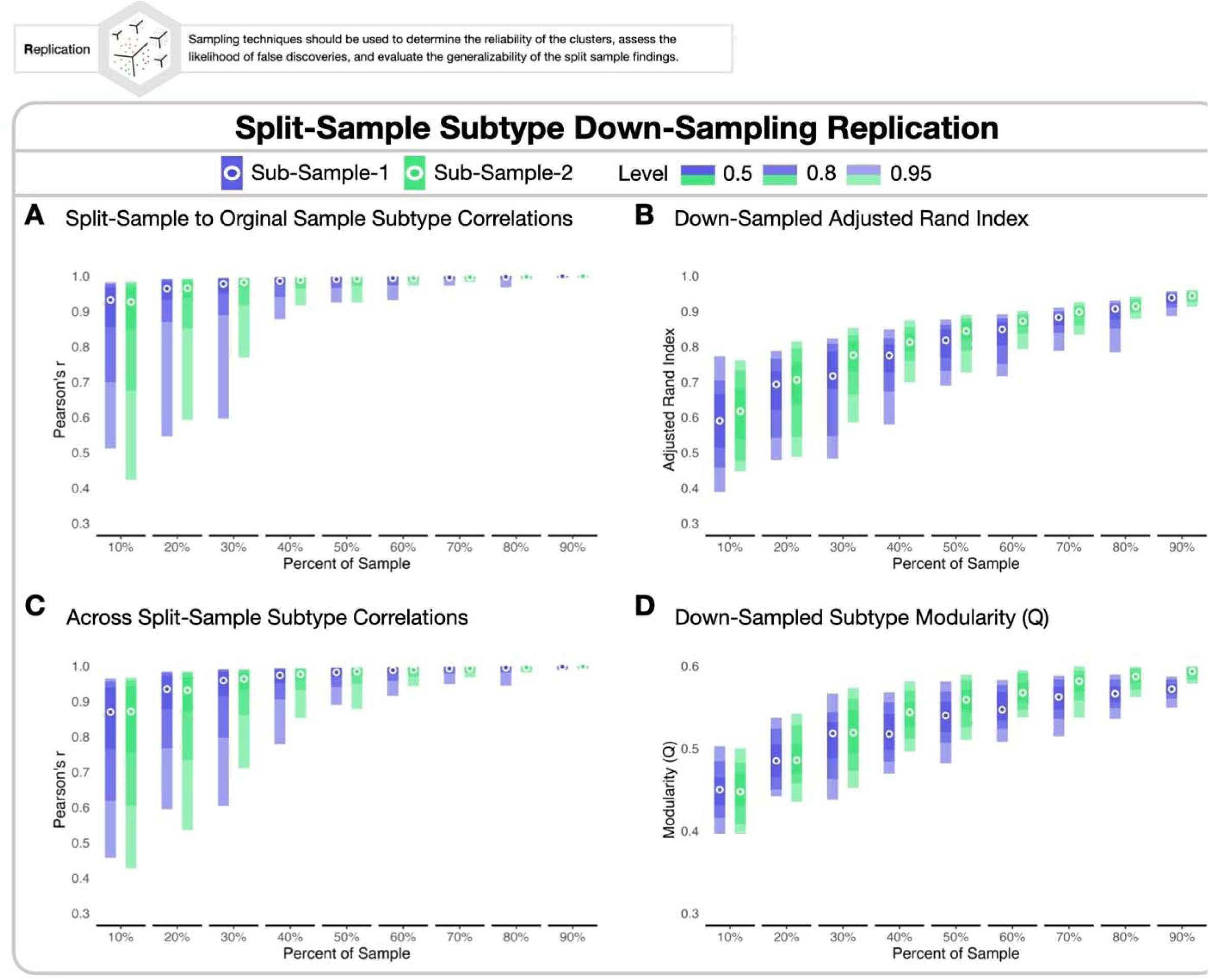
Split-sample Subtype Down-Sampling Replication. For each increment of down-sampling, ranging from 10% to 90%. Shaded bars indicate 95% (lightest shading), .8% (mid shading), and .5 (darkest shading) confidence interval and are colored by sample (Sub-Sample-1; blue, Sub-Sample-2; green). **A)** Mean maximum correlations with the down-samples to the original samples by split. **B)** Adjusted rand index between down-samples and original samples. **C)** Mean maximum correlations across split samples. **D)** Down-sampled final bagged modularity (Q) by sample.

## Discussion

### The Subtypes and their associations to cognition and mental health

The current study set out with three primary goals: to identify subtypes of children based on their RSFC, to explore the potential of these subtypes to act as brain-based predictors for cognitive abilities and mental health, and to investigate the reproducibility and reliability of these subtypes. We identified four distinct neurodevelopmental subtypes using cortical and subcortical RSFC connections that offer insight into the relationship between RSFC, demographics, cognitive functioning, and mental health. Importantly, these subtypes were highly reproducible across different samples and were not solely influenced by socioeconomic or demographic factors. However, these factors did differ across some of the subtypes, underscoring the potential influence of environmental factors, such as socio-economic status (Moriguchi & Shinohara, 2019), parental education (Dubow et al., 2009), and adversity (McLaughlin et al., 2014; Wade et al., 2022), as being associated with brain connectivity patterns. These patterns, in turn, seem to be associated with specific cognitive and mental health outcomes. First, we discuss the implications of our subtypes concerning their RSFC profiles and phenotypic relationships. Then, we discuss how the IVEPR framework enhanced the reliability of our conclusions, which posit the subtypes as neuro-markers for cognitive and mental health in children and adolescents. Finally, we discuss the broader implications of our findings.

The distinct connectivity profiles within the subtypes offer a window into heterogeneous whole-brain functional profiles underlying cognitive functioning and mental health in late-grade-school children. Subtype-1 and Subtype-4 are marked by functional connectivity patterns that appear to be associated with higher cognitive functioning levels and lower mental health problems. These two subtypes share some common features in their connectivity profiles. The strong connectivity within the default mode network may support higher degrees of internally based thought, including potential evaluation and introspection (Luo et al., 2016; Zhang et al., 2022). Such an internal focus may facilitate cognitive processes like abstract reasoning and planning, which may potentially be fostered by the socioeconomically advantaged backgrounds that were found to be associated with this subtype (Aartsen et al., 2019). The strong negative connectivity between auditory and sensorimotor-hand networks may indicate a separation of sensory inputs from motor outputs, potentially leading to more refined motor control and sensory discrimination and better cognitive performance (Gordon et al., 2023).

In addition, the negative connectivity between the default mode and dorsal attention network aligns with typical anticorrelation between these networks during rest (Dixon et al., 2016; Owens et al., 2020). This finding indicates a conventional pattern of brain connectivity that may underlie efficient cognitive processing and attentional control. However, other aspects of the connectivity profiles between these two subtypes are distinct and emphasize the concept of nested heterogeneity, suggesting that similar cognitive and mental health outcomes may arise from different underlying RSFC patterns. For example, Subtype 1 has strong negative connectivity between the default mode network and the cingulo-opercular network, while Subtype 4 has positive connectivity between these networks.

Similarly, Subtypes 2 and 3, both of which are associated with greater degrees of cognitive and mental health difficulties than Subtypes 1 and 2, also share some similar aspects of their connectivity profiles that differentiate them from Subtypes 1 and 2. In particular, they exhibit negative connectivity within the default mode network, within the cingulo-opercular network, and between the default mode network and regions that do not fall into any organized network (i.e., none). In contrast, Subtypes 1 and 4 show positive connectivity for these connections. In addition, Subtypes 2 and 3 show positive connectivity between the default mode network and the dorsal attention network, which contrasts with the negative connectivity for this aspect of connectivity shown for Subtypes 1 and 4.

Yet once again, nested heterogeneity is evident in the connectivity profiles distinguishing Subtypes 2 and 3, each characterized by unique patterns of lower-order sensorimotor connectivity. Subtype-2 is characterized by the strongest negative connectivity within and between the sensorimotor networks, indicating a form of sensorimotor integration that might be less conducive to efficient cognitive processing. Conversely, Subtype-3 distinguishes itself with strong positive connections between auditory and sensorimotor networks, suggesting a different mode of sensorimotor coordination that may support more effective rapid response mechanisms in specific contexts (Adise et al., 2022; Karcher & Barch, 2021). These distinct connectivity configurations within lower-order networks underscore the diverse ways sensorimotor integration can influence cognitive performance across these subtypes. At present, the reason these pattern configurations relate to lower cognitive performance is a subject of speculation, and we are actively investigating this question. Notably, Subtypes 1 and 4, which exhibited higher cognitive performance and fewer mental health issues, showed connectivity in higher-order systems such as the default mode and cingulo-opercular networks.

In summary, our RSFC subtypes provide insight into the distinct brain connectivity patterns associated with cognitive functioning and mental health in children and adolescents. Subtypes 1 and 4, marked by specific connectivity configurations of integration between higher-order networks (e.g., dorsal attention, default mode network), are associated with higher cognitive abilities and fewer mental health issues. In contrast, Subtypes 2 and 3 are characterized by a contrasting connectivity pattern that may favor immediate sensory-motor responses over higher-order cognitive processing. The degree to which these connectivity patterns might be linked to environmental and socio-economic factors must be investigated in future studies.

### The IVEPR framework

The IVEPR framework was a fundamental component in achieving the aims of this study, as it provided a comprehensive toolset for identifying, evaluating, and validating the RSFC subtypes. The framework’s efficacy was demonstrated through the split-sample approach, which confirmed that the identified RSFC subtypes were highly reproducible. The ability to replicate these subtypes even with a down-sample as small as 10% of the original size is particularly noteworthy, as it suggests that the subtypes are fundamentally stable and can be reliably identified across different sample populations and sizes. This implies that the subtypes are not just artifacts of a particular dataset but may reflect underlying individual differences in their functional brain architecture. The reliability of these subtypes lays the groundwork for future studies. It suggests that subsequent research can build on these findings to investigate neurodevelopmental patterns and behavioral characteristics that may be associated with them. Establishing which phenotypic measures consistently differentiate these subtypes across the two sub-samples allowed us to add another layer of validation to the subtypes. This step was critical for understanding which measures reliably differed and reproduced between the subtypes across the two sub-samples (i.e., Sub-Sample-1, Sub-Sample-2).

Perhaps the most significant implication of the IVEPR framework’s application is the demonstration that the RSFC subtypes have the most consistent predictive value for cognitive functioning and mental health profiles. This finding supported our hypothesis that an individual’s whole functional brain profile may offer more insight into cognitive and emotional functioning profiles than isolated RSFC connections. Our findings highlight the diverse roles different brain regions or networks might have in shaping developmental pattern differences. When discussing ’different brain profiles,’ we refer to unique connectivity patterns within the brain, as identified through RSFC data. We show that despite varying connectivity profiles, two subtypes are associated with higher cognitive performance and fewer mental health issues, while two other subtypes exhibit lower cognitive performance and greater mental health challenges. This suggests that there may not be optimal or adverse functional connectivity configuration for cognitive and mental health outcomes. Instead, multiple configurations can lead to similar cognitive or mental health patterns, indicating that certain brain regions or networks may have a more pronounced impact on specific cognitive processes or mental health conditions than previously understood. It will be necessary for future studies to assess if these subtypes have practical relevance in tracking developmental progress over time and informing clinical interventions.

The reproducible identification of RSFC subtypes across individuals who meet typical inclusion criteria and those often excluded from neurodevelopmental research due to noisy data could have significant implications for improving retention and representation in these studies. Historically, individuals who exhibit high motion during scans, often those with pronounced behavioral problems or cognitive impairments (Thomson et al., 2021), are excluded from neurodevelopmental research. This exclusion has limited our understanding of brain-behavior relationships in those who might benefit most from such insights (Satterthwaite et al., 2012). However, our findings indicate that even those typically excluded due to noisy data may still be reliably subtyped. Notably, the subtype profiles of this high-motion sample showed robust reproducibility to those in Sub-Sample-1 and Sub-Sample-2 of our primary analyses. In examining the cognitive functioning and mental health differences across Sub-Sample-1 and Sub-Sample-2 in the “complete sample,” we identified significant differences in attention problems, accuracy on the emotional Stroop task, and executive-capability-(BPPCA) among the subtypes that were not replicated across Sub-Sample-1 and Sub-Sample-2 of our primary analyses. The crucial point with this finding is that including high-motion individuals in the analysis allowed us to identify additional cognitive functioning and mental health differences between the subtypes. These findings suggest that our subtyping approach may allow future studies to include those previously excluded, which could ultimately enhance our understanding of brain-behavior relationships. Future studies will need to investigate this idea in more detail.

## Limitations and Future Directions

While this study offers valuable insights into the heterogeneity of RSFC subtypes and their association with cognitive functioning and mental health, several limitations warrant consideration. Our study exclusively analyzed RSFC data, limiting our understanding to brain connectivity patterns without considering structural differences or task-based activations, which might provide broader insights into brain metrics across subtypes. Future research incorporating multi-modal imaging, including diffusion tensor imaging (DTI), structural MRI, and task-based fMRI, could enhance our understanding of these subtypes and their generalizability, potentially offering a more comprehensive perspective on brain neuromarkers (Calhoun & Sui, 2016; Ooi et al., 2022), and the utility of a multi-modal approach in subtype analysis. Finally, our study’s cross-sectional design limits our ability to infer causal relationships between RSFC subtypes and developmental outcomes. Longitudinal data will be necessary to elucidate these associations’ directionality and determine whether these subtypes predict changes in cognitive and mental health outcomes over time. We are currently examining this issue.

The findings from our study pave the way for several future research avenues. First, a deeper exploration into the longitudinal development and stability of these RSFC subtypes is warranted. Understanding how these subtypes evolve through late childhood/adolescence and the factors influencing their stability or change may provide important insights into neurodevelopment. Such a longitudinal approach may illuminate whether these subtypes are transient phases or stable markers of brain organization throughout an individual’s life. Finally, considering the extensive range of measures gathered in the ABCD study, there is considerable opportunity to delineate further brain-behavior relationships between these RSFC subtypes. Investigating these additional relationships with these measures would be worthwhile in obtaining a more comprehensive characterization of these RSFC subtypes.

## Conclusions

Our study represents a significant step in parsing heterogeneous patterns of brain organization based on an individual’s resting-state functional whole-brain profile to be used to predict cognitive functioning and mental health during late childhood. Through the IVEPR framework, we successfully identified and validated four distinct RSFC subtypes and demonstrated their robustness and reliability across diverse sample sizes. These results suggest that the RSFC subtypes are reliable neuro-markers for tracking variations amongst individuals in their functional neural organization. In addition, this study sets a benchmark for future studies to build off using these RSFC subtypes to investigate how they influence the developmental trajectories of each subtype. In addition, these findings underscore the potential of subtypes as pivotal tools in neuroscientific research. Exploring further applications and potential uses of these subtypes and the IVEPR framework in future studies is likely to be useful.

## Acknowledgements

Data used in the preparation of this manuscript were obtained from the Adolescent Brain Cognitive Development (ABCD) Study (https://abcdstudy.org), held in the NIMH Data Archive (NDA). The ABCD Study® is supported by the National Institutes of Health, and additional federal partners under Award nos. U01DA041048, U01DA050989, U01DA051016, U01DA041022, U01DA051018, U01DA051037, U01DA050987, U01DA041174, U01DA041106, U01DA041117, U01DA041028, U01DA041134, U01DA050988, U01DA051039, U01DA041156, U01DA041025, U01DA041120, U01DA051038, U01DA041148, U01DA041093, U01DA041089, U24DA041123, U24DA041147. A full list of supporters is available at https://abcdstudy.org/federal-partners.html. A listing of participating sites and a complete listing of the study investigators can be found at https://abcdstudy.org/consortium_members/. ABCD consortium investigators designed and implemented the study and/or provided data but did not necessarily participate in the analysis or writing of this report.

