## Supplemental Tables for "Neurodevelopmental Subtypes of Functional Brain Organization in the ABCD Study Using a Rigorous Analytic Framework": Neurodevelopmental Subtypes Supplemental Tables.docx

| **Supplemental Table 1** | | | | | | | | | | | | | | | | |
| --- | --- | --- | --- | --- | --- | --- | --- | --- | --- | --- | --- | --- | --- | --- | --- | --- |
|  | | **Full-Sample** | | | | | **Sub-Sample-1** | | | | | **Sub-Sample-2** | | | | |
|  |  | **Sample** | **Subtype 1** | **Subtype 2** | **Subtype 3** | **Subtype 4** | **Sample** | **Subtype 1** | **Subtype 2** | **Subtype 3** | **Subtype 4** | **Sample** | **Subtype 1** | **Subtype 2** | **Subtype 3** | **Subtype 4** |
|  |  | N=7293 | N=2062 (28.27%) | N=1776 (24.35%) | N=1645 (22.56%) | N=1810 (24.82%) | N=3670 | N=984 (26.81%) | N=928 (25.29%) | N=805 (21.93%) | N=953 (25.97%) | N=3623 | N=1027 (28.35%) | N=813 (22.44%) | N=874 (24.12%) | N=909 (25.09%) |
| **Age** | **F (p) or X2 (p)** |  | 0.82 | | | |  | 0.8 | | | |  | 2.68* | | | |
|  | **Post-Hoc** |  |  |  |  |  |  |  |  |  |  |  |  |  |  |  |
|  | **Mean (SD) or N (%)** | 9.93 (0.62) | 9.92 (0.62) | 9.91 (0.61) | 9.95 (0.62) | 9.92 (0.61) | 9.93 (0.62) | 9.92 (0.61) | 9.95 (0.61) | 9.94 (0.63) | 9.91 (0.62) | 9.92 (0.61) | 9.93 (0.62) | 9.87 (0.60) | 9.94 (0.62) | 9.93 (0.61) |
| **Sex at Birth** |  |  | 3.27 | | | |  | 3.5 | | | |  | 1.5 | | | |
| Female | |  |  |  |  |  |  |  |  |  |  |  |  |  |  |  |
|  |  | 3608 (49.47%) | 1030 (49.95%) | 846 (47.64%) | 829 (50.4%) | 903 (49.89%) | 1849 (50.38%) | 506 (51.42%) | 446 (48.06%) | 420 (52.17%) | 477 (50.05%) | 1759 (48.55%) | 487 (47.42%) | 388 (47.72%) | 433 (49.54%) | 451 (49.61%) |
| Male | |  |  |  |  |  |  |  |  |  |  |  |  |  |  |  |
|  |  | 3685 (50.53%) | 1032 (50.05%) | 930 (52.36%) | 816 (49.6%) | 907 (50.11%) | 1821 (49.62%) | 478 (48.58%) | 482 (51.94%) | 385 (47.83%) | 476 (49.95%) | 1864 (51.45%) | 540 (52.58%) | 425 (52.28%) | 441 (50.46%) | 458 (50.39%) |
| **Parent Education** |  |  | 420.47*** | | | |  | 233.72*** | | | |  | 216.19*** | | | |
| < HS Diploma |  |  | < 3, > 4 | < 3, > 4 | > 1, > 2, > 4 | < 1, < 2, < 3 |  | < 3 | < 3, > 4 | > 1, > 2, > 4 | < 2, < 3 |  | < 3, > 4 | < 3 | > 1, > 2, > 4 | < 1, < 3 |
|  |  | 340 (4.67%) | 80 (3.89%) | 75 (4.23%) | 143 (8.69%) | 42 (2.32%) | 183 (4.99%) | 35 (3.56%) | 47 (5.09%) | 76 (9.44%) | 25 (2.62%) | 157 (4.34%) | 39 (3.8%) | 28 (3.44%) | 72 (8.24%) | 18 (1.98%) |
| HS Diploma/GED | |  | < 2, < 3 | > 1, < 3, > 4 | > 1, > 2, > 4 | < 2, < 3 |  | < 3, > 4 | < 3, > 4 | > 1, > 2, > 4 | < 1, < 2, < 3 |  | < 2, < 3 | > 1, < 3 | > 1, > 2, > 4 | < 3 |
|  |  | 632 (8.67%) | 122 (5.93%) | 157 (8.86%) | 247 (15.02%) | 106 (5.86%) | 298 (8.13%) | 65 (6.61%) | 71 (7.68%) | 122 (15.16%) | 40 (4.2%) | 334 (9.22%) | 60 (5.85%) | 69 (8.49%) | 134 (15.33%) | 71 (7.81%) |
| Some College | |  | < 3 | < 3 | > 1, > 2, > 4 | < 3 |  | < 3 | < 3 | > 1, > 2, > 4 | < 3 |  | < 3 | < 3 | > 1, > 2, > 4 | < 3 |
|  |  | 1863 (25.57%) | 459 (22.29%) | 424 (23.93%) | 579 (35.2%) | 401 (22.15%) | 920 (25.1%) | 214 (21.77%) | 231 (25.0%) | 273 (33.91%) | 202 (21.2%) | 943 (26.04%) | 228 (22.24%) | 198 (24.35%) | 313 (35.81%) | 204 (22.44%) |
| Bachelor | |  | > 3 | > 3 | < 1, < 2, < 4 | > 3 |  | > 2, > 3 | < 1, > 3, < 4 | < 1, < 2, < 4 | > 2, > 3 |  | > 3 | > 3 | < 1, < 2, < 4 | > 3 |
|  |  | 1850 (25.39% | 567 (27.54%) | 458 (25.85%) | 305 (18.54%) | 520 (28.73%) | 948 (25.87%) | 278 (28.28%) | 223 (24.13%) | 161 (20.0%) | 286 (30.01%) | 902 (24.91%) | 276 (26.93%) | 215 (26.45%) | 157 (17.96%) | 254 (27.94%) |
| Post Graduate Degree | |  | > 2, > 3 | < 1, > 3, < 4 | > 1, < 2, < 4 | > 2, > 3 |  | > 3 | > 3 | < 1, < 2, < 4 | > 3 |  | > 3 | > 3 | < 1, < 2, < 4 | > 3 |
|  |  | 2601 (35.7%) | 831 (40.36%) | 658 (37.13%) | 371 (22.55%) | 741 (40.94%) | 1316 (35.91%) | 391 (39.78%) | 352 (38.1%) | 173 (21.49%) | 400 (41.97%) | 1285 (35.49%) | 422 (41.17%) | 303 (37.27%) | 198 (22.65%) | 362 (39.82%) |
| **Household Income** |  |  | 368.04*** | | | |  | 217.29*** | | | |  | 166.57*** | | | |
| <50K | |  | < 2, < 3, > 4 | > 1, < 3, > 4 | > 1, > 2, > 4 | < 1, < 2, < 3 |  | < 2, < 3 | > 1, < 3, > 4 | > 1, > 2, > 4 | < 2, < 3 |  | < 3 | < 3 | > 1, > 2, > 4 | < 3 |
|  |  | 1895 (28.37%) | 431 (22.65%) | 450 (27.51%) | 677 (46.92%) | 337 (19.85%) | 930 (27.68%) | 191 (21.1%) | 248 (28.67%) | 333 (47.37%) | 158 (17.81%) | 965 (29.07%) | 229 (24.03%) | 193 (25.91%) | 357 (46.54%) | 186 (21.75%) |
| >=50K & <100K | |  | < 4 | < 4 | < 4 | > 1, > 2, > 3 |  |  | < 4 | < 4 | > 2, > 3 |  |  |  |  |  |
|  |  | 1893 (28.34%) | 532 (27.96%) | 451 (27.57%) | 378 (26.2%) | 532 (31.33%) | 978 (29.11%) | 266 (29.39%) | 231 (26.71%) | 189 (26.88%) | 292 (32.92%) | 915 (27.56%) | 257 (26.97%) | 206 (27.65%) | 197 (25.68%) | 255 (29.82%) |
| >=100K | |  | > 2, > 3 | < 1, > 3, < 4 | < 1, < 2, < 4 | > 2, > 3 |  | > 2, > 3 | < 1, > 3 | < 1, < 2, < 4 | > 3 |  | > 3 | > 3 | < 1, < 2, < 4 | > 3 |
|  |  | 2892 (43.29%) | 940 (49.4%) | 735 (44.93%) | 388 (26.89%) | 829 (48.82%) | 1452 (43.21%) | 448 (49.5%) | 386 (44.62%) | 181 (25.75%) | 437 (49.27%) | 1440 (43.37%) | 467 (49.0%) | 346 (46.44%) | 213 (27.77%) | 414 (48.42%) |
| **Parental Marital Status** |  |  | 335.11*** | | | |  | 199.45*** | | |  |  | 154.22*** | | | |
| No | |  | < 2, < 3, > 4 | > 1, < 3, > 4 | > 1, > 2, > 4 | < 1, < 2, < 3 |  | < 2, < 3 | > 1, < 3, > 4 | > 1, > 2, > 4 | < 2, < 3 |  | < 3, > 4 | < 3, > 4 | > 1, > 2, > 4 | < 1, < 2, < 3 |
|  |  | 2228 (30.79%) | 517 (25.27%) | 540 (30.56%) | 783 (48.3%) | 388 (21.53%) | 1115 (30.56%) | 238 (24.34%) | 280 (30.34%) | 397 (49.94%) | 200 (21.01%) | 1113 (31.02%) | 269 (26.42%) | 235 (29.05%) | 409 (47.56%) | 200 (22.2%) |
| Yes | |  | > 2, > 3, > 4 | < 1, > 3, < 4 | < 1, < 2, < 4 | < 1, > 2, > 3 |  | > 2, > 3 | < 1, > 3, < 4 | < 1, < 2, < 4 | > 2, > 3 |  | > 3, > 4 | > 3, < 4 | < 1, < 2, < 3 | < 1, > 2, > 3 |
|  |  | 5008 (69.21%) | 1529 (74.73%) | 1227 (69.44%) | 838 (51.7%) | 1414 (78.47%) | 2533 (69.44%) | 740 (75.66%) | 643 (69.66%) | 398 (50.06%) | 752 (78.99%) | 2475 (68.98%) | 749 (73.58%) | 574 (70.95%) | 451 (52.44%) | 701 (77.8%) |
| **Supplemental Table 1.** "passed RSFC quality control" Sample Distribution of Demographic and Socioeconomic Variables across the Full-Sample, Sub-Sample-1, and Sub-Sample-2. Significant p-values are denoted as *p<0.05, **p<0.01, ***p<0.001. Significant FDR post-hoc comparisons are indicated by < or > between Subtypes. | | | | | | | | | | | | | | | | |

| **Supplemental Table 2** | | | | | | | | | |
| --- | --- | --- | --- | --- | --- | --- | --- | --- | --- |
| **Domain** | **Measure** | **Sample** | **Subtype 1** | **Subtype 2** | **Subtype 3** | **Subtype 4** | **F** | **Post Hoc** | **Robust** |
|  | | | **M (SD)** | **M (SD)** | **M (SD)** | **M (SD)** |  | | |
| **Adversity** | **adversity** | Full-Sample | 0.35 (0.92) | 0.39 (1.00) | 0.59 (1.17) | 0.30 (0.88) | 28.37*** | 1=2<3>4 |  |
|  |  | Sub-Sample-1 | 0.34 (0.91) | 0.40 (1.03) | 0.64 (1.24) | 0.29 (0.90) | 18.21*** | 1=2<3>4 | TRUE |
|  |  | Sub-Sample-2 | 0.35 (0.94) | 0.38 (0.98) | 0.56 (1.09) | 0.29 (0.83) | 12.93*** | 1=2<3>4 | TRUE |
| **Psychopathology (CBCL)** | **aggressive** | Full-Sample | -0.32 (3.84) | -0.08 (4.21) | 0.24 (4.71) | -0.29 (3.85) | 6.83*** | 3>1=4 |  |
|  |  | Sub-Sample-1 | -0.42 (3.64) | -0.21 (4.08) | 0.40 (4.76) | -0.25 (4.07) | 6.41*** | 3>1=2=4 |  |
|  |  | Sub-Sample-2 | -0.18 (4.12) | -0.09 (4.17) | 0.11 (4.66) | -0.30 (3.66) | 1.59 |  |  |
|  | **anxious depressed** | Full-Sample | 0.02 (2.29) | -0.01 (2.31) | -0.09 (2.26) | 0.06 (2.38) | 1.29 |  |  |
|  |  | Sub-Sample-1 | -0.12 (2.13) | -0.02 (2.29) | 0.02 (2.35) | 0.13 (2.49) | 1.92 |  |  |
|  |  | Sub-Sample-2 | 0.13 (2.41) | -0.04 (2.27) | -0.18 (2.16) | 0.03 (2.35) | 2.97* | 1>3 |  |
|  | **attention problems** | Full-Sample | -0.39 (3.09) | 0.16 (3.57) | -0.06 (3.35) | -0.07 (3.39) | 8.98*** | 2>1, 3=4>1 |  |
|  |  | Sub-Sample-1 | -0.44 (2.99) | 0.16 (3.55) | 0.08 (3.47) | -0.16 (3.38) | 6.41*** | 2=3>1 | TRUE |
|  |  | Sub-Sample-2 | -0.33 (3.24) | 0.10 (3.44) | -0.15 (3.28) | 0.03 (3.43) | 3.04* | 2>1 | TRUE |
|  | **externalizing** | Full-Sample | -0.17 (1.70) | -0.04 (1.90) | 0.14 (2.14) | -0.15 (1.69) | 10.77*** | 3>2>1, 3>4 |  |
|  |  | Sub-Sample-1 | -0.21 (1.62) | -0.07 (1.82) | 0.20 (2.16) | -0.12 (1.79) | 7.97*** | 3>2>1, 3>4 | TRUE |
|  |  | Sub-Sample-2 | -0.12 (1.84) | -0.06 (1.87) | 0.11 (2.12) | -0.16 (1.59) | 3.62* | 3>1=4 | TRUE |
|  | **internalizing** | Full-Sample | -0.02 (1.57) | -0.01 (1.65) | 0.00 (1.66) | 0.00 (1.58) | 0.06 |  |  |
|  |  | Sub-Sample-1 | -0.09 (1.45) | -0.01 (1.66) | 0.05 (1.73) | 0.03 (1.66) | 1.5 |  |  |
|  |  | Sub-Sample-2 | 0.04 (1.63) | -0.04 (1.58) | -0.03 (1.62) | 0.02 (1.57) | 0.6 |  |  |
|  | **rule breaking** | Full-Sample | -0.19 (1.60) | -0.03 (1.83) | 0.20 (2.05) | -0.15 (1.56) | 17.7*** | 1=4<2<3 |  |
|  |  | Sub-Sample-1 | -0.21 (1.57) | -0.02 (1.76) | 0.20 (2.05) | -0.12 (1.67) | 9*** | 1=4<2<3 | TRUE |
|  |  | Sub-Sample-2 | -0.16 (1.73) | -0.10 (1.77) | 0.21 (2.05) | -0.18 (1.45) | 9.85*** | 1=2<3, 4 < 3 | TRUE |
|  | **social problems** | Full-Sample | -0.17 (2.06) | 0.00 (2.31) | 0.06 (2.31) | -0.17 (2.12) | 5.24*** | 1=2<3, 4 < 3 |  |
|  |  | Sub-Sample-1 | -0.30 (1.90) | -0.02 (2.34) | 0.11 (2.35) | -0.13 (2.23) | 5.57*** | 1>2, 3>1 |  |
|  |  | Sub-Sample-2 | 0.02 (2.31) | -0.06 (2.15) | 0.02 (2.22) | -0.20 (2.04) | 2.04 |  |  |
|  | **somatic complaints** | Full-Sample | 0.01 (1.90) | -0.00 (1.97) | 0.04 (2.06) | 0.01 (1.91) | 0.13 |  |  |
|  |  | Sub-Sample-1 | -0.02 (1.81) | 0.03 (1.99) | 0.02 (2.12) | -0.03 (1.88) | 0.2 |  |  |
|  |  | Sub-Sample-2 | 0.02 (1.95) | -0.06 (1.92) | 0.07 (2.03) | 0.07 (1.96) | 0.83 |  |  |
|  | **thought problems** | Full-Sample | -0.13 (1.95) | 0.09 (2.26) | -0.06 (2.23) | 0.02 (2.08) | 3.98** | 2>1 |  |
|  |  | Sub-Sample-1 | -0.22 (1.87) | 0.05 (2.20) | 0.01 (2.32) | -0.01 (2.04) | 3.12* | 1=2<3, 1<4 |  |
|  |  | Sub-Sample-2 | -0.04 (2.07) | 0.09 (2.24) | -0.10 (2.15) | 0.06 (2.15) | 1.37 |  |  |
|  | **total problems** | Full-Sample | -0.18 (1.72) | 0.04 (1.97) | 0.02 (1.98) | -0.07 (1.80) | 5.48*** | 3>1=2 |  |
|  |  | Sub-Sample-1 | -0.25 (1.61) | 0.02 (1.95) | 0.09 (2.03) | -0.08 (1.87) | 6.04*** | 3>1=2 |  |
|  |  | Sub-Sample-2 | -0.08 (1.88) | 0.00 (1.88) | -0.03 (1.93) | -0.05 (1.78) | 0.36 |  |  |
|  | **withdrawn depressed** | Full-Sample | -0.08 (1.62) | -0.01 (1.69) | 0.05 (1.75) | -0.06 (1.59) | 2.26 |  |  |
|  |  | Sub-Sample-1 | -0.15 (1.51) | -0.03 (1.67) | 0.12 (1.78) | -0.02 (1.66) | 3.92** | 3>1 |  |
|  |  | Sub-Sample-2 | -0.03 (1.67) | -0.04 (1.67) | 0.01 (1.75) | -0.06 (1.56) | 0.24 |  |  |
| **Cognitive and EF** | **LMT** | Full-Sample | 0.03 (0.17) | -0.00 (0.17) | -0.01 (0.16) | 0.02 (0.16) | 26.57*** | 1=4>2>3 |  |
|  |  | Sub-Sample-1 | 0.04 (0.17) | -0.00 (0.17) | -0.01 (0.16) | 0.02 (0.17) | 14.55*** | 1=4 >2=3 | TRUE |
|  |  | Sub-Sample-2 | 0.02 (0.16) | 0.01 (0.17) | -0.02 (0.16) | 0.02 (0.17) | 10.39*** | 1=4>2>3 | TRUE |
|  | **RAVLT** | Full-Sample | 1.45 (9.67) | 0.33 (9.83) | -1.32 (9.54) | 1.65 (9.34) | 33.19*** | 1=4>2>3 |  |
|  |  | Sub-Sample-1 | 1.63 (9.46) | 0.54 (9.90) | -1.13 (9.59) | 1.93 (9.69) | 16.96*** | 1=4>2>3 | TRUE |
|  |  | Sub-Sample-2 | 1.21 (9.79) | 0.39 (9.95) | -1.51 (9.39) | 1.24 (9.03) | 15.35*** | 1=2=4>3 | TRUE |
|  | **general-ability-(BPPCA)** | Full-Sample | 0.17 (0.70) | 0.07 (0.74) | -0.19 (0.74) | 0.17 (0.71) | 96.33*** | 1=4>2>3 |  |
|  |  | Sub-Sample-1 | 0.19 (0.68) | 0.07 (0.75) | -0.18 (0.75) | 0.19 (0.70) | 50.4*** | 1=4>2>3 | TRUE |
|  |  | Sub-Sample-2 | 0.14 (0.71) | 0.09 (0.72) | -0.20 (0.72) | 0.17 (0.71) | 48.97*** | 1=2>3, 4>2>3 | TRUE |
|  | **executive-capability-(BPPCA)** | Full-Sample | 0.08 (0.72) | 0.01 (0.73) | -0.06 (0.78) | 0.07 (0.70) | 12.73*** | 1=4>2>3 |  |
|  |  | Sub-Sample-1 | 0.08 (0.72) | 0.01 (0.74) | -0.09 (0.79) | 0.07 (0.71) | 8.81*** | 1=2=4>3 | TRUE |
|  |  | Sub-Sample-2 | 0.08 (0.73) | 0.01 (0.72) | -0.02 (0.76) | 0.07 (0.69) | 4.2** | 1=4>3 | TRUE |
|  | **learning/memory-(BPPCA)** | Full-Sample | 0.11 (0.69) | 0.02 (0.71) | -0.11 (0.67) | 0.11 (0.67) | 37.61*** | 1=4>2>3 |  |
|  |  | Sub-Sample-1 | 0.12 (0.68) | 0.03 (0.70) | -0.10 (0.67) | 0.14 (0.67) | 20.68*** | 1=4>2>3 | TRUE |
|  |  | Sub-Sample-2 | 0.10 (0.69) | 0.04 (0.71) | -0.13 (0.68) | 0.06 (0.66) | 18.55*** | 1=2=4>3 | TRUE |
|  | **common-EF-(CFA)** | Full-Sample | 0.14 (0.75) | 0.01 (0.81) | -0.18 (0.87) | 0.13 (0.73) | 60.42*** | 1=4>2>3 |  |
|  |  | Sub-Sample-1 | 0.15 (0.73) | 0.02 (0.85) | -0.22 (0.90) | 0.14 (0.74) | 36.18*** | 1=4>2>3 | TRUE |
|  |  | Sub-Sample-2 | 0.13 (0.77) | 0.01 (0.75) | -0.16 (0.84) | 0.13 (0.73) | 25.87*** | 1=4>2>3 | TRUE |
|  | **cognitive-aptitude-(CFA)** | Full-Sample | 0.17 (0.76) | 0.04 (0.85) | -0.26 (0.87) | 0.17 (0.74) | 101.86*** | 1=4>2>3 |  |
|  |  | Sub-Sample-1 | 0.20 (0.75) | 0.05 (0.86) | -0.26 (0.88) | 0.18 (0.74) | 57.78*** | 1=4>2>3 | TRUE |
|  |  | Sub-Sample-2 | 0.13 (0.79) | 0.04 (0.81) | -0.27 (0.87) | 0.15 (0.75) | 48.66*** | 1=4>2>3 | TRUE |
|  | **updating-specific-(CFA)** | Full-Sample | 0.10 (0.67) | 0.04 (0.69) | -0.19 (0.71) | 0.11 (0.64) | 66.83*** | 1=4>2>3 |  |
|  |  | Sub-Sample-1 | 0.14 (0.66) | 0.05 (0.68) | -0.16 (0.70) | 0.12 (0.64) | 35.13*** | 1=4>2>3 | TRUE |
|  |  | Sub-Sample-2 | 0.07 (0.68) | 0.05 (0.68) | -0.22 (0.71) | 0.10 (0.64) | 36.54*** | 1=2>3, 4>3 | TRUE |
| **Stroop** | **angry acc eq** | Full-Sample | 0.01 (0.07) | 0.00 (0.07) | -0.00 (0.08) | 0.01 (0.06) | 6.61*** | 1>2=3, 4>2>3 |  |
|  |  | Sub-Sample-1 | 0.01 (0.06) | 0.00 (0.07) | -0.00 (0.08) | 0.00 (0.07) | 2.98* | 1>3 | TRUE |
|  |  | Sub-Sample-2 | 0.01 (0.07) | 0.00 (0.07) | -0.00 (0.07) | 0.01 (0.06) | 4.32** | 1>3, 4>2=3 | TRUE |
|  | **happy acc eq** | Full-Sample | 0.01 (0.07) | 0.00 (0.07) | -0.00 (0.08) | 0.01 (0.07) | 6.23*** | 1>3, 4>2=3 |  |
|  |  | Sub-Sample-1 | 0.01 (0.06) | 0.00 (0.07) | -0.00 (0.08) | 0.01 (0.07) | 3.73* | 1>2=3 |  |
|  |  | Sub-Sample-2 | 0.01 (0.07) | 0.00 (0.07) | -0.00 (0.08) | 0.01 (0.07) | 2.96* |  |  |
|  | **stroop interference** | Full-Sample | 0.01 (0.06) | -0.00 (0.06) | -0.00 (0.07) | 0.00 (0.06) | 5.35*** | 1>2=3, 4>3 |  |
|  |  | Sub-Sample-1 | 0.01 (0.06) | -0.00 (0.06) | -0.00 (0.08) | 0.00 (0.06) | 3.6* | 1>2=3 |  |
|  |  | Sub-Sample-2 | 0.01 (0.06) | -0.00 (0.06) | 0.00 (0.07) | 0.00 (0.05) | 1.88 |  |  |
| **UPPS** | **negative urgency** | Full-Sample | -0.04 (0.42) | -0.00 (0.43) | 0.03 (0.45) | -0.01 (0.41) | 7.47*** | 3 >1>2=4, |  |
|  |  | Sub-Sample-1 | -0.04 (0.41) | -0.02 (0.43) | 0.01 (0.44) | -0.02 (0.42) | 2.57 |  |  |
|  |  | Sub-Sample-2 | -0.03 (0.43) | -0.00 (0.43) | 0.05 (0.46) | 0.01 (0.41) | 5.49*** | 3=4>1, 3>2 |  |
|  | **perseverance** | Full-Sample | -0.02 (0.31) | 0.01 (0.32) | -0.01 (0.34) | 0.00 (0.32) | 3.82** | 2>1 |  |
|  |  | Sub-Sample-1 | -0.02 (0.31) | 0.01 (0.32) | -0.02 (0.33) | -0.00 (0.32) | 2.27 |  |  |
|  |  | Sub-Sample-2 | -0.02 (0.31) | -0.00 (0.31) | -0.00 (0.35) | 0.01 (0.31) | 1.09 |  |  |
|  | **positive urgency** | Full-Sample | -0.05 (0.49) | 0.00 (0.50) | 0.05 (0.52) | -0.02 (0.48) | 15.18*** | 3 >2>1, 1>4, 3>4 |  |
|  |  | Sub-Sample-1 | -0.07 (0.48) | -0.01 (0.51) | 0.04 (0.51) | -0.02 (0.48) | 7*** | 1 < 2 < 3, 4 < 1, 1 < 4, 4 < 3 | TRUE |
|  |  | Sub-Sample-2 | -0.04 (0.49) | -0.01 (0.50) | 0.07 (0.54) | -0.01 (0.48) | 8.45*** | 1=2<3, 4<3 | TRUE |
|  | **premeditation** | Full-Sample | -0.02 (0.34) | 0.01 (0.35) | -0.01 (0.37) | -0.00 (0.33) | 3.16* | 2 >1 |  |
|  |  | Sub-Sample-1 | -0.02 (0.34) | 0.02 (0.35) | -0.01 (0.36) | -0.01 (0.33) | 2.26 |  |  |
|  |  | Sub-Sample-2 | -0.01 (0.34) | -0.00 (0.35) | 0.00 (0.38) | 0.00 (0.32) | 0.42 |  |  |
|  | **sensation seeking** | Full-Sample | -0.01 (0.41) | 0.01 (0.40) | 0.00 (0.41) | 0.02 (0.40) | 1.7 |  |  |
|  |  | Sub-Sample-1 | -0.02 (0.40) | 0.01 (0.40) | 0.00 (0.42) | 0.00 (0.40) | 0.92 |  |  |
|  |  | Sub-Sample-2 | 0.00 (0.41) | 0.01 (0.40) | 0.01 (0.42) | 0.02 (0.39) | 0.28 |  |  |
| **Supplemental Table 2.** “passed RSFC quality control” sample ANOVA comparisons across various phenotypic domains across the Full-Sample, Sub-Sample-1, and Sub-Sample-2. For each domain, the table lists mean (M) and standard deviation (SD) values for four subtypes. The 'F' column shows the ANOVA results with significant p-values are denoted as *p<0.05, **p<0.01, ***p<0.001, 'Post Hoc' column shows the FDR-corrected post-hoc comparisons between the subtypes. The 'Robust' column indicates if at least one of the subtypes post-hoc comparisons was replicated across Sub-Sample-1 and Sub-Sample-2. | | | | | | | | | |

| **Supplemental Table 3** | | | | | | | | | | | | | | | | |
| --- | --- | --- | --- | --- | --- | --- | --- | --- | --- | --- | --- | --- | --- | --- | --- | --- |
|  | | **Full Sample** | | | | | **Sample-1** | | | | | **Sample  2** | | | | |
|  |  | **Sample** | **Subtype 1** | **Subtype 2** | **Subtype 3** | **Subtype 4** | **Sample** | **Subtype 1** | **Subtype 2** | **Subtype 3** | **Subtype 4** | **Sample** | **Subtype 1** | **Subtype 2** | **Subtype 3** | **Subtype 4** |
|  |  | N=9027 | N=2531 (28.04%) | N=2503 (27.73%) | N=2081 (23.05%) | N=1912 (21.18%) | N=4513 | N=1231 (27.28%) | N=1229 (27.23%) | N=1114 (24.68%) | N=939 (20.81%) | N=4514 | N=1243 (27.54%) | N=1243 (27.54%) | N=1015 (22.49%) | N=1013 (22.44%) |
| **Age** | **F (p) or X2 (p)** |  | 2.32 |  |  |  |  | 0.48 |  |  |  |  | 1.54 |  |  |  |
|  | **Post-Hoc** |  |  |  |  |  |  |  |  |  |  |  |  |  |  |  |
|  | **Mean (SD) or N (%)** | 9.91 (0.62) | 9.91 (0.62) | 9.89 (0.61) | 9.94 (0.63) | 9.90 (0.61) | 9.91 (0.62) | 9.91 (0.61) | 9.90 (0.62) | 9.93 (0.63) | 9.90 (0.62) | 9.9 (0.61) | 9.90 (0.62) | 9.87 (0.61) | 9.93 (0.62) | 9.91 (0.61) |
| **Sex at Birth** |  |  | 3.54 |  |  |  |  | 3.35 |  |  |  |  | 4.51 |  |  |  |
| Female |  |  |  |  |  |  |  |  |  |  |  |  |  |  |  |  |
|  |  | 4281 (47.42%) | 1209 (47.77%) | 1153 (46.06%) | 934 (48.85%) | 985 (47.33%) | 2162 (47.91%) | 601 (48.9%) | 566 (45.98%) | 465 (49.52%) | 530 (47.58%) | 2119 (46.94%) | 570 (45.86%) | 562 (45.21%) | 495 (48.77%) | 492 (48.57%) |
| Male |  |  |  |  |  |  |  |  |  |  |  |  |  |  |  |  |
|  |  | 4746 (52.58%) | 1322 (52.23%) | 1350 (53.94%) | 978 (51.15%) | 1096 (52.67%) | 2351 (52.09%) | 628 (51.1%) | 665 (54.02%) | 474 (50.48%) | 584 (52.42%) | 2395 (53.06%) | 673 (54.14%) | 681 (54.79%) | 520 (51.23%) | 521 (51.43%) |
| **Parent Education** |  |  | 496.79*** | | |  |  | 261.68*** | | |  |  | 227.23*** | | | |
| < HS Diploma |  |  | < 3, > 4 | < 3, > 4 | > 1, > 2, > 4 | < 1, < 2, < 3 |  | < 3 | < 3, > 4 | > 1, > 2, > 4 | < 2, < 3 |  | < 3, > 4 | < 3, > 4 | > 1, > 2, > 4 | < 1,  2, < 3 |
|  |  | 452 (5.01%) | 103 (4.08%) | 115 (4.61%) | 179 (9.36%) | 55 (2.64%) | 237 (5.26%) | 50 (4.08%) | 63 (5.13%) | 90 (9.58%) | 34 (3.05%) | 215 (4.77%) | 53 (4.27%) | 48 (3.86%) | 90 (8.87%) | 24 (2.37%) |
| HS Diploma/GED |  |  | < 2, < 3 | > 1, < 3, > 4 | > 1, > 2, > 4 | < 2, < 3 |  | < 2, < 3 | > 1, < 3, > 4 | > 1, > 2, > 4 | < 2, < 3 |  | < 2, < 3 | > 1, < 3 | > 1, > 2, > 4 | < 3 |
|  |  | 856 (9.49%) | 165 (6.53%) | 246 (9.85%) | 310 (16.21%) | 135 (6.49%) | 405 (8.99%) | 82 (6.69%) | 117 (9.54%) | 150 (15.97%) | 56 (5.03%) | 451 (10.0%) | 84 (6.77%) | 118 (9.5%) | 164 (16.16%) | 85 (8.39%) |
| Some College |  |  | < 2, < 3 | > 1, < 3 | < 1, > 2, > 4 | < 3 |  | < 3 | < 3 | > 1, > 2, > 4 | < 3 |  | < 3 | < 3 | > 1, > 2, > 4 | < 3 |
|  |  | 2366 (26.24%) | 577 (22.83%) | 634 (25.39%) | 673 (35.2%) | 482 (23.16%) | 1159 (25.72%) | 276 (22.51%) | 310 (25.26%) | 325 (34.61%) | 248 (22.26%) | 1207 (26.76%) | 284 (22.88%) | 324 (26.09%) | 358 (35.27%) | 241 (23.79%) |
| Bachelor |  |  | > 3 | > 3 | < 1, < 2, < 4 | > 3 |  | > 3 | > 3, < 4 | < 1, < 2, < 4 | > 2, > 3 |  | > 3 | > 3 | < 1, < 2, < 4 | > 3 |
|  |  | 2234 (24.78%) | 679 (26.87%) | 630 (25.23%) | 347 (18.15%) | 578 (27.78%) | 1153 (25.59%) | 336 (27.41%) | 301 (24.53%) | 187 (19.91%) | 329 (29.53%) | 1081 (23.96%) | 327 (26.35%) | 304 (24.48%) | 175 (17.24%) | 275 (27.15%) |
| Post Graduate Degree |  |  | > 2, > 3 | < 1, > 3, > 4 | > 1, < 2, < 4 | < 2, > 3 |  | > 3 | > 3, < 4 | < 1, < 2, < 4 | > 2, > 3 |  | > 3 | > 3 | < 1, < 2, < 4 | > 3 |
|  |  | 3109 (34.48%) | 1003 (39.69%) | 872 (34.92%) | 403 (21.08%) | 831 (39.93%) | 1552 (34.44%) | 482 (39.31%) | 436 (35.53%) | 187 (19.91%) | 447 (40.13%) | 1557 (34.52%) | 493 (39.73%) | 448 (36.07%) | 228 (22.46%) | 388 (38.3%) |
| **Household Income** |  |  | 451.44*** | | |  |  | 246.59*** | | |  |  | 196.31*** | | | |
| <50K |  |  | < 2, < 3 | > 1, < 3, > 4 | > 1, > 2, > 4 | < 2, < 3 |  | < 2, < 3 | > 1, < 3, > 4 | > 1, > 2, > 4 | > 2, > 3 |  | < 3 | < 3, > 4 | > 1, > 2, > 4 | < 2, < 3 |
|  |  | 2473 (29.95%) | 555 (23.79%) | 676 (29.44%) | 826 (49.25%) | 416 (21.33%) | 1225 (29.68%) | 259 (22.92%) | 352 (30.88%) | 405 (49.27%) | 209 (20.17%) | 1248 (30.23%) | 287 (24.96%) | 311 (27.43%) | 430 (48.21%) | 220 (23.11%) |
| >=50K & <100K |  |  |  | > 3 | < 2, < 4 | > 3 |  |  |  | < 4 | > 3 |  |  |  | < 4 | > 3 |
|  |  | 2341 (28.36%) | 651 (27.9%) | 659 (28.7%) | 434 (25.88%) | 597 (30.62%) | 1191 (28.85%) | 325 (28.76%) | 317 (27.81%) | 222 (27.01%) | 327 (31.56%) | 1150 (27.86%) | 317 (27.57%) | 320 (28.22%) | 228 (25.56%) | 285 (29.94%) |
| >=100K |  |  | > 2, > 3 | < 1, > 3, > 4 | < 1, < 2, < 4 | < 2, > 3 |  | > 2, > 3 | < 1, > 3, < 4 | < 1, < 2, < 4 | > 2, > 3 |  | > 3 | > 3 | < 1, < 2 | > 3 |
|  |  | 3442 (41.69%) | 1127 (48.31%) | 961 (41.86%) | 417 (24.87%) | 937 (48.05%) | 1712 (41.47%) | 546 (48.32%) | 471 (41.32%) | 195 (23.72%) | 500 (48.26%) | 1730 (41.91%) | 546 (47.48%) | 503 (44.36%) | 234 (26.23%) | 447 (46.95%) |
| **Parental Marital Status** |  |  | 383.23*** | | |  |  | 235.93*** | | |  |  | 152.57*** | | | |
| No |  |  | < 2, < 3, > 4 | > 1, < 3, > 4 | > 1, > 2, > 4 | < 1, < 2, < 3 |  | < 2, < 3 | > 1, < 3, > 4 | > 1, > 2, > 4 | < 2, < 3 |  | < 2, < 3 | > 1, < 3, > 4 | > 1, > 2, > 4 | < 2, < 3 |
|  |  | 2917 (32.58%) | 670 (26.68%) | 814 (32.74%) | 946 (50.19%) | 487 (23.52%) | 1448 (32.29%) | 314 (25.72%) | 401 (32.73%) | 482 (51.94%) | 251 (22.59%) | 1469 (32.88%) | 341 (27.68%) | 399 (32.33%) | 481 (48.24%) | 248 (24.68%) |
| Yes |  |  | > 2, > 3, > 4 | < 1, > 3, > 4 | < 1, < 2, < 4 | < 1, < 2, > 3 |  | > 2, > 3 | < 1, > 3, < 4 | < 1, < 2, < 4 | > 2, > 3 |  | > 2, > 3 | < 1, > 3, > 4 | < 1, < 2, < 4 | < 2, > 3 |
|  |  | 6036 (67.42%) | 1841 (73.32%) | 1672 (67.26%) | 939 (49.81%) | 1584 (76.48%) | 3037 (67.71%) | 907 (74.28%) | 824 (67.27%) | 446 (48.06%) | 860 (77.41%) | 2999 (67.12%) | 891 (72.32%) | 835 (67.67%) | 516 (51.76%) | 757 (75.32%) |
| **Supplemental Table 3.**  "Complete" Sample Distribution of Demographic and Socioeconomic Variables across the Full Sample, Sample-1, and Sample-2. Significant p-values are denoted as *p<0.05, **p<0.01, ***p<0.001. Significant FDR post-hoc comparisons are indicated by < or > between Subtypes. | | | | | | | | | | | | | | | | |

| **Supplemental Table 4** | | | | | | |
| --- | --- | --- | --- | --- | --- | --- |
|  | | **Full Sample** | | | | |
|  |  | **Sample** | **Subtype 1** | **Subtype 2** | **Subtype 3** | **Subtype 4** |
|  |  | N=1293 | N=306 (23.67%) | N=312 (24.13%) | N=232 (17.94%) | N=443 (34.26%) |
| **Age** | **F or X2 (p)** |  | 0.9 |  |  |  |
|  | **Post-Hoc** |  |  |  |  |  |
|  | **Mean (SD) or N (%)** | 9.8 (0.6) | 9.75 (0.57) | 9.81 (0.61) | 9.81 (0.61) | 9.82 (0.59) |
| **Sex at Birth** |  |  | 5.69 |  |  |  |
| Female |  |  | > 2, < 4 | < 1 |  | > 1 |
|  |  | 466 (36.04%) | 127 (41.5%) | 104 (33.33%) | 84 (36.21%) | 151 (34.09%) |
| Male |  |  | < 2, < 4 | > 1 |  | > 1 |
|  |  | 827 (63.96%) | 179 (58.5%) | 208 (66.67%) | 148 (63.79%) | 292 (65.91%) |
| **Parent Education** |  |  | 52.05*** | | | |
| < HS Diploma |  |  |  | < 3 | > 2, > 4 | < 3 |
|  |  | 87 (6.74%) | 21 (6.89%) | 16 (5.14%) | 27 (11.64%) | 23 (5.2%) |
| HS Diploma/GED |  |  | < 3 | < 3 | > 1, > 2, > 4 | < 3 |
|  |  | 163 (12.64%) | 39 (12.79%) | 38 (12.22%) | 44 (18.97%) | 42 (9.5%) |
| Some College |  |  | < 3 | < 3 | > 1, > 2, < 4 | > 3 |
|  |  | 379 (29.38%) | 84 (27.54%) | 84 (27.01%) | 84 (36.21%) | 127 (28.73%) |
| Bachelor |  |  |  |  |  |  |
|  |  | 283 (21.94%) | 64 (20.98%) | 78 (25.08%) | 44 (18.97%) | 97 (21.95%) |
| Post Graduate Degree |  |  | > 3 | > 3 | < 1, < 2, < 4 | > 3 |
|  |  | 378 (29.3%) | 97 (31.8%) | 95 (30.55%) | 33 (14.22%) | 153 (34.62%) |
| **Household Income** |  |  | 65.34*** | | | |
| <50K |  |  | < 3 | < 3 | > 1, > 2, > 4 | < 3 |
|  |  | 445 (37.81%) | 98 (35.0%) | 97 (34.52%) | 126 (60.87%) | 124 (30.32%) |
| >=50K & <100K |  |  |  |  |  |  |
|  |  | 323 (27.44%) | 73 (26.07%) | 85 (30.25%) | 47 (22.71%) | 118 (28.85%) |
| >=100K |  |  | > 3 | > 3 | < 1, < 2, < 4 | > 3 |
|  |  | 409 (34.75%) | 109 (38.93%) | 99 (35.23%) | 34 (16.43%) | 167 (40.83%) |
| **Parental Marital Status** |  |  | 34.62*** | | | |
| No |  |  | < 3 | < 3 | > 1, > 2, < 4 | > 3 |
|  |  | 517 (40.36%) | 115 (37.7%) | 118 (38.06%) | 131 (57.46%) | 153 (34.93%) |
| Yes |  |  | > 3 | > 3 | < 1, < 2 , < 4 | > 3 |
|  |  | 764 (59.64%) | 190 (62.3%) | 192 (61.94%) | 97 (42.54%) | 285 (65.07%) |
| **Supplemental Table 4**."High Motion" Sample Distribution of Demographic and Socioeconomic Variables across the Full Sample, Sample-1, and Sample-2. Significant p-values are denoted as *p<0.05, **p<0.01, ***p<0.001. Significant FDR post-hoc comparisons are indicated by < or > between Subtypes. | | | | | | |

| **Supplement Table 5** | | | | | | | | | | |
| --- | --- | --- | --- | --- | --- | --- | --- | --- | --- | --- |
| **Data** | **Sub-Sample** | **Model** | **Df** | **AIC** | **BIC** | **Chisq** | **Chisq diff** | **RMSEA** | **Df diff** | **P Chisq.** |
| **UPPS** | Sample-1 | Configural | 492 | 161232.539 | 162870.789 | 1215.677 |  |  |  |  |
|  | Sample-2 |  | 492 | 159803.477 | 161438.535 | 1299.751 |  |  |  |  |
|  | Sample-1 | Metric | 537 | 161228.801 | 162587.803 | 1301.939 | 86.262 | 0.032 | 45 | 0 |
|  | Sample-2 |  | 537 | 159765.728 | 161122.083 | 1352.002 | 52.251 | 0.013 | 45 | 0.213 |
|  | Sample-1 | Scalar | 576 | 161238.198 | 162355.187 | 1389.337 | 87.398 | 0.037 | 39 | 0 |
|  | Sample-2 |  | 576 | 159760.4 | 160875.212 | 1424.674 | 72.672 | 0.031 | 39 | 0.001 |
|  | Sample-1 | Strict | 630 | 161334.111 | 162116.003 | 1593.25 | 203.913 | 0.055 | 54 | 0 |
|  | Sample-2 |  | 630 | 159830.796 | 160611.165 | 1603.071 | 178.396 | 0.05 | 54 | 0 |
|  | Sample-1 | Means | 645 | 161345.278 | 162034.088 | 1634.417 | 41.167 | 0.044 | 15 | 0 |
|  | Sample-2 |  | 645 | 159847.278 | 160534.746 | 1649.553 | 46.482 | 0.048 | 15 | 0 |
| **Data** | **Sub-Sample** | **Model** | **Df** | **AIC** | **BIC** | **Chisq** | **Chisq diff** | **RMSEA** | **Df diff** | **P Chisq.** |
| **Cognitive and EF** | Sample-1 | Configural | 40 | 161681.963 | 162293.096 | 72.789 |  |  |  |  |
|  | Sample-2 |  | 40 | 157722.788 | 158331.614 | 71.446 |  |  |  |  |
|  | Sample-1 | Metric | 58 | 161668.931 | 162170.06 | 95.757 | 22.968 | 0.018 | 18 | 0.192 |
|  | Sample-2 |  | 58 | 157701.311 | 158200.548 | 85.969 | 14.523 | 0 | 18 | 0.694 |
|  | Sample-1 | Scalar | 70 | 161692.458 | 162120.251 | 143.284 | 47.527 | 0.06 | 12 | 0 |
|  | Sample-2 |  | 70 | 157740.709 | 158166.887 | 149.367 | 63.397 | 0.073 | 12 | 0 |
|  | Sample-1 | Strict | 91 | 161813.033 | 162112.488 | 305.859 | 162.575 | 0.09 | 21 | 0 |
|  | Sample-2 |  | 91 | 157828.012 | 158126.336 | 278.67 | 129.303 | 0.08 | 21 | 0 |
|  | Sample-1 | Means | 100 | 161944.703 | 162189.156 | 455.529 | 149.67 | 0.137 | 9 | 0 |
|  | Sample-2 |  | 100 | 157957.354 | 158200.884 | 426.012 | 147.342 | 0.137 | 9 | 0 |
| Measurement Invariance of the UPPS Impulsive Behavior Scale and the Cognitive and Executive Function (EF) across two independent samples. Shown are the degrees of freedom (Df), Akaike Information Criterion (AIC), Bayesian Information Criterion (BIC), Chi-square (Chisq), Chi-square difference (Chisq diff), Root Mean Square Error of Approximation (RMSEA), degrees of freedom difference (Df diff), and the p-value of the Chi-square test (P Chisq). Each model was examined to assess the model fit, the invariance across groups, and the robustness of the measures, aiding in understanding the underlying structure and comparative fit of the impulsivity and cognitive/executive function constructs across different samples. | | | | | | | | | | |
| **Sub-Sample-1 Confirmatory Factor Analysis Cognitive and EF Model Loadings** | | | | | | |  | | | |
| **Latent Variables** | **Estimate** | **Std.Err** | **z-value** | **P(>\|z\|)** | **Std.lv** | **Std.all** |  |  |  |  |
| **CommonEF** |  |  |  |  |  |  |  |  |  |  |
| CardSort | 6.93 | 0.19 | 36 | 0 | 6.93 | 0.751 |  |  |  |  |
| Flanker | 5.146 | 0.167 | 31 | 0 | 5.146 | 0.579 |  |  |  |  |
| nb | 6.062 | 0.27 | 22 | 0 | 6.062 | 0.432 |  |  |  |  |
| List | 5.037 | 0.222 | 23 | 0 | 5.037 | 0.433 |  |  |  |  |
| sst | -0.482 | 0.159 | -3 | 0.002 | -0.482 | -0.057 |  |  |  |  |
| **UpdatingSpecific** |  |  |  |  |  |  |  |  |  |  |
| nb | 4.493 | 0.339 | 13 | 0 | 4.493 | 0.32 |  |  |  |  |
| List_r | 4.909 | 0.319 | 15 | 0 | 4.909 | 0.422 |  |  |  |  |
| **Cognitive Ability** |  |  |  |  |  |  |  |  |  |  |
| matrix | 2.395 | 0.068 | 35 | 0 | 2.395 | 0.638 |  |  |  |  |
| PicVocab | 4.791 | 0.139 | 34 | 0 | 4.791 | 0.614 |  |  |  |  |
| **Sub-Sample-2 Confirmatory Factor Analysis Cognitive and EF Model Loadings** | | | | | | |  |  |  |  |
| **Latent Variables** | **Estimate** | **Std.Err** | **z-value** | **P(>\|z\|)** | **Std.lv** | **Std.all** |  |  |  |  |
| **CommonEF** |  |  |  |  |  |  |  |  |  |  |
| CardSort | 5.915 | 0.178 | 33 | 0 | 5.915 | 0.669 |  |  |  |  |
| Flanker | 5.296 | 0.168 | 32 | 0 | 5.296 | 0.614 |  |  |  |  |
| nb | 6.188 | 0.28 | 22 | 0 | 6.188 | 0.437 |  |  |  |  |
| List | 5.503 | 0.234 | 23 | 0 | 5.503 | 0.463 |  |  |  |  |
| sst | -0.954 | 0.157 | -6 | 0 | -0.954 | -0.118 |  |  |  |  |
| **UpdatingSpecific** |  |  |  |  |  |  |  |  |  |  |
| nb | 4.21 | 0.343 | 12 | 0 | 4.21 | 0.297 |  |  |  |  |
| List | 5.249 | 0.347 | 15 | 0 | 5.249 | 0.442 |  |  |  |  |
| **Cognitive Ability** |  |  |  |  |  |  |  |  |  |  |
| matrix | 2.321 | 0.068 | 34 | 0 | 2.321 | 0.622 |  |  |  |  |
| PicVocab | 4.791 | 0.142 | 34 | 0 | 4.791 | 0.614 |  |  |  |  |
| **Sub-Sample-1 Confirmatory Factor Analysis UPPS Model Loadings** | | | | | | |  |  |  |  |
| **Latent Variables** | **Estimate** | **Std.Err** | **z-value** | **P(>\|z\|)** | **Std.lv** | **Std.all** |  |  |  |  |
| **premeditation** |  |  |  |  |  |  |  |  |  |  |
| upps6_y | 0.469 | 0.016 | 30 | 0 | 0.469 | 0.631 |  |  |  |  |
| upps16 | 0.424 | 0.016 | 27 | 0 | 0.424 | 0.531 |  |  |  |  |
| upps21_y | -0.177 | 0.023 | -8 | 0 | -0.177 | -0.165 |  |  |  |  |
| upps27_y | -0.096 | 0.025 | -4 | 0 | -0.096 | -0.082 |  |  |  |  |
| **perseverance** |  |  |  |  |  |  |  |  |  |  |
| upps15_y | 0.382 | 0.013 | 30 | 0 | 0.382 | 0.499 |  |  |  |  |
| upps19_y | 0.535 | 0.013 | 40 | 0 | 0.535 | 0.641 |  |  |  |  |
| upps22_y | 0.541 | 0.012 | 45 | 0 | 0.541 | 0.711 |  |  |  |  |
| upps24_y | 0.398 | 0.012 | 33 | 0 | 0.398 | 0.538 |  |  |  |  |
| **sensation seeking** |  |  |  |  |  |  |  |  |  |  |
| upps12_y | 0.581 | 0.024 | 25 | 0 | 0.581 | 0.555 |  |  |  |  |
| upps18_y | 0.415 | 0.02 | 21 | 0 | 0.415 | 0.442 |  |  |  |  |
| upps21_y | 0.319 | 0.024 | 13 | 0 | 0.319 | 0.297 |  |  |  |  |
| upps27_y | 0.566 | 0.027 | 21 | 0 | 0.566 | 0.483 |  |  |  |  |
| **negative urgency** |  |  |  |  |  |  |  |  |  |  |
| upps7_y | 0.502 | 0.018 | 28 | 0 | 0.502 | 0.486 |  |  |  |  |
| upps11_y | 0.438 | 0.014 | 32 | 0 | 0.438 | 0.541 |  |  |  |  |
| upps17_y | 0.538 | 0.017 | 32 | 0 | 0.538 | 0.537 |  |  |  |  |
| upps20_y | 0.577 | 0.016 | 36 | 0 | 0.577 | 0.607 |  |  |  |  |
| **positive urgency** |  |  |  |  |  |  |  |  |  |  |
| upps35_y | 0.551 | 0.013 | 43 | 0 | 0.551 | 0.64 |  |  |  |  |
| upps36_y | 0.659 | 0.015 | 44 | 0 | 0.659 | 0.65 |  |  |  |  |
| upps37_y | 0.707 | 0.013 | 54 | 0 | 0.707 | 0.772 |  |  |  |  |
| upps39_y | 0.654 | 0.015 | 44 | 0 | 0.654 | 0.655 |  |  |  |  |
| **Sub-Sample-2 Confirmatory Factor Analysis UPPS Model Loadings** | | | | | | |  |  |  |  |
| **Latent Variables** | **Estimate** | **Std.Err** | **z-value** | **P(>\|z\|)** | **Std.lv** | **Std.all** |  |  |  |  |
| **premeditation** |  |  |  |  |  |  |  |  |  |  |
| upps6_y | 0.469 | 0.016 | 30 | 0 | 0.469 | 0.631 |  |  |  |  |
| upps16 | 0.424 | 0.016 | 27 | 0 | 0.424 | 0.531 |  |  |  |  |
| upps21_y | -0.177 | 0.023 | -8 | 0 | -0.177 | -0.165 |  |  |  |  |
| upps27_y | -0.096 | 0.025 | -4 | 0 | -0.096 | -0.082 |  |  |  |  |
| **perseverance** |  |  |  |  |  |  |  |  |  |  |
| upps15_y | 0.382 | 0.013 | 30 | 0 | 0.382 | 0.499 |  |  |  |  |
| upps19_y | 0.535 | 0.013 | 40 | 0 | 0.535 | 0.641 |  |  |  |  |
| upps22_y | 0.541 | 0.012 | 45 | 0 | 0.541 | 0.711 |  |  |  |  |
| upps24_y | 0.398 | 0.012 | 33 | 0 | 0.398 | 0.538 |  |  |  |  |
| **sensation seeking** |  |  |  |  |  |  |  |  |  |  |
| upps12_y | 0.581 | 0.024 | 25 | 0 | 0.581 | 0.555 |  |  |  |  |
| upps18_y | 0.415 | 0.02 | 21 | 0 | 0.415 | 0.442 |  |  |  |  |
| upps21_y | 0.319 | 0.024 | 13 | 0 | 0.319 | 0.297 |  |  |  |  |
| upps27_y | 0.566 | 0.027 | 21 | 0 | 0.566 | 0.483 |  |  |  |  |
| **negative urgency** |  |  |  |  |  |  |  |  |  |  |
| upps7_y | 0.502 | 0.018 | 28 | 0 | 0.502 | 0.486 |  |  |  |  |
| upps11_y | 0.438 | 0.014 | 32 | 0 | 0.438 | 0.541 |  |  |  |  |
| upps17_y | 0.538 | 0.017 | 32 | 0 | 0.538 | 0.537 |  |  |  |  |
| upps20_y | 0.577 | 0.016 | 36 | 0 | 0.577 | 0.607 |  |  |  |  |
| **positive urgency** |  |  |  |  |  |  |  |  |  |  |
| upps35_y | 0.551 | 0.013 | 43 | 0 | 0.551 | 0.64 |  |  |  |  |
| upps36_y | 0.659 | 0.015 | 44 | 0 | 0.659 | 0.65 |  |  |  |  |
| upps37_y | 0.707 | 0.013 | 54 | 0 | 0.707 | 0.772 |  |  |  |  |
| upps39_y | 0.654 | 0.015 | 44 | 0 | 0.654 | 0.655 |  |  |  |  |

| **Supplement Table 6** | | | | | | |
| --- | --- | --- | --- | --- | --- | --- |
| **RSFC Metric** | **Top 1** | **Top 5** | **Top 10** | **Subcortical ROI legend** | **Subcortical ROI** | **Subcortical ROI Name** |
| Subtype | 5 (18.52%) | 11 (40.74%) | 17 (62.96%) |  | crcxlh | left-cerebellum-cortex |
| dt-dla | 4 (14.81%) | 7 (25.93%) | 11 (40.74%) |  | aglh | left-amygdala |
| ca-ca | 3 (11.11%) | 8 (29.63%) | 10 (37.04%) |  | crcxrh | right-cerebellum-cortex |
| dla-crcxlh | 2 (7.41%) | 5 (18.52%) | 8 (29.63%) |  | vtdclh | left-ventral diencephalon |
| smh-aglh | 2 (7.41%) | 3 (11.11%) |  |  | cdelh | left-caudate |
| vta-crcxrh | 2 (7.41%) | 3 (11.11%) |  |  | hplh | left-hippocampus |
| ad-rspltp | 1 (3.7%) |  |  |  | aarh | right-accumbens-area |
| ca-aglh | 1 (3.7%) | 2 (7.41%) |  |  | thplh | left-thalamus-proper |
| ca-fo | 1 (3.7%) | 2 (7.41%) |  |  | vtdcrh | right-ventral diencephalon |
| cgc-vtdclh | 1 (3.7%) | 7 (25.93%) | 11 (40.74%) |  | pllh | left-pallidum |
| dla-cdelh | 1 (3.7%) |  |  |  |  |  |
| dla-hplh | 1 (3.7%) |  | 4 (14.81%) |  |  |  |
| dt-fo | 1 (3.7%) |  |  |  |  |  |
| smh-aarh | 1 (3.7%) | 3 (11.11%) | 4 (14.81%) |  |  |  |
| smm-aglh | 1 (3.7%) |  |  |  |  |  |
| cgc-thplh |  | 4 (14.81%) | 8 (29.63%) |  |  |  |
| dt-dt |  | 4 (14.81%) | 4 (14.81%) |  |  |  |
| dla-n |  | 3 (11.11%) | 5 (18.52%) |  |  |  |
| vta-vtdcrh |  | 3 (11.11%) |  |  |  |  |
| ad-ad |  | 2 (7.41%) | 4 (14.81%) |  |  |  |
| cgc-aglh |  |  | 5 (18.52%) |  |  |  |
| smh-pllh |  |  | 5 (18.52%) |  |  |  |
| fo-crcxlh |  |  | 4 (14.81%) |  |  |  |
| sa-vtdclh |  |  | 4 (14.81%) |  |  |  |
| **Supplemental Table 6.** Top Resting State Functional Connectivity (RSFC) metrics in predicting cognitive functioning and mental health measures across the Conditional Random Field (CRF) models. The table quantifies the frequency and proportion of each RSFC metric emerging as a top predictor in the models. Each row represents a distinct RSFC metric along with its occurrence and percentage in the top 1, 5, and 10 ranks across the models, illuminating their predictive prowess in the examined cognitive and mental health domains. | | | | | | |

| **Supplement Table 7** | | | | | |
| --- | --- | --- | --- | --- | --- |
| **Sample** | **Domain** | **Measure** | **Full Sample** | **Sample-1** | **Sample-2** |
| **Include Only** | **CBCL** | aggressive | F=4.72 (1.58), p=0.03 (0.06) | F=4.42 (1.55), p=0.04 (0.07) | F=1.38 (0.8), p=0.5 (0.25) |
|  |  | anxious depressed | F=1.2 (0.74), p=0.47 (0.26) | F=1.55 (0.8), p=0.33 (0.24) | F=2.24 (0.96), p=0.3 (0.19) |
|  |  | attention problems | F=6.05 (1.67), p=0.01 (0.03) | F=4.4 (1.43), p=0.04 (0.06) | F=2.33 (1.07), p=0.3 (0.21) |
|  |  | externalizing | F=7.21 (1.98), p=0 (0.02) | F=5.41 (1.73), p=0.03 (0.06) | F=2.66 (1.19), p=0.26 (0.22) |
|  |  | internalizing | F=0.45 (0.37), p=0.77 (0.2) | F=1.32 (0.71), p=0.39 (0.25) | F=0.75 (0.54), p=0.7 (0.2) |
|  |  | rule breaking | F=11.6 (2.52), p=0 (0) | F=6.1 (1.79), p=0.02 (0.04) | F=6.58 (1.92), p=0.02 (0.04) |
|  |  | social problems | F=3.7 (1.36), p=0.06 (0.1) | F=3.91 (1.29), p=0.05 (0.08) | F=1.64 (0.81), p=0.43 (0.22) |
|  |  | somatic complaints | F=0.47 (0.38), p=0.76 (0.2) | F=0.52 (0.42), p=0.71 (0.22) | F=0.89 (0.58), p=0.65 (0.22) |
|  |  | thought problems | F=2.95 (1.16), p=0.11 (0.13) | F=2.37 (0.98), p=0.17 (0.16) | F=1.27 (0.75), p=0.54 (0.23) |
|  |  | total problems | F=3.89 (1.33), p=0.05 (0.09) | F=4.2 (1.36), p=0.04 (0.07) | F=0.62 (0.46), p=0.73 (0.2) |
|  |  | withdrawn depressed | F=1.84 (1), p=0.29 (0.24) | F=2.85 (1.15), p=0.12 (0.13) | F=0.53 (0.4), p=0.77 (0.18) |
|  | **Cognitive Functioning** | LMT | F=17.12 (2.86), p=0 (0) | F=9.61 (2.16), p=0 (0) | F=6.89 (1.77), p=0 (0.01) |
|  |  | RAVLT | F=21.33 (3.19), p=0 (0) | F=11.16 (2.46), p=0 (0) | F=10.08 (2.17), p=0 (0) |
|  |  | general-capability-(BPPCA) | F=61.5 (5.64), p=0 (0) | F=32.34 (4.26), p=0 (0) | F=31.25 (3.96), p=0 (0) |
|  |  | executive-capability-(BPPCA) | F=8.5 (2.08), p=0 (0) | F=5.84 (1.76), p=0 (0.02) | F=3.06 (1.21), p=0.07 (0.1) |
|  |  | learning/memory-(BPPCA) | F=24.06 (3.4), p=0 (0) | F=13.53 (2.67), p=0 (0) | F=12.12 (2.4), p=0 (0) |
|  |  | common-EF-(CFA) | F=38.68 (4.37), p=0 (0) | F=23.18 (3.58), p=0 (0) | F=16.79 (2.98), p=0 (0) |
|  |  | cognitive-aptitude-(CFA) | F=64.97 (5.78), p=0 (0) | F=36.98 (4.53), p=0 (0) | F=31.22 (4.12), p=0 (0) |
|  |  | updating specific-(CFA) | F=42.74 (4.71), p=0 (0) | F=22.76 (3.5), p=0 (0) | F=23.5 (3.55), p=0 (0) |
|  | **Stroop** | angry acc eq | F=4.54 (1.49), p=0.02 (0.04) | F=2.26 (1.01), p=0.17 (0.16) | F=3.06 (1.2), p=0.13 (0.16) |
|  |  | happy acc eq | F=4.3 (1.45), p=0.03 (0.04) | F=2.75 (1.09), p=0.13 (0.13) | F=2.18 (1.01), p=0.21 (0.19) |
|  |  | stroop interference | F=3.74 (1.32), p=0.04 (0.06) | F=2.6 (1.02), p=0.13 (0.13) | F=1.53 (0.81), p=0.31 (0.22) |
|  | **UPPS** | negative urgency | F=5.16 (1.57), p=0.02 (0.03) | F=2 (0.91), p=0.26 (0.19) | F=3.87 (1.41), p=0.07 (0.12) |
|  |  | perseverance | F=2.78 (1.12), p=0.12 (0.14) | F=1.82 (0.83), p=0.29 (0.2) | F=1.06 (0.63), p=0.58 (0.26) |
|  |  | positive urgency | F=10.05 (2.21), p=0 (0) | F=4.81 (1.49), p=0.05 (0.09) | F=5.76 (1.75), p=0.02 (0.05) |
|  |  | premeditation | F=2.38 (1.06), p=0.16 (0.16) | F=1.84 (0.89), p=0.29 (0.21) | F=0.65 (0.47), p=0.69 (0.23) |
|  |  | sensation seeking | F=1.47 (0.8), p=0.32 (0.23) | F=0.93 (0.59), p=0.52 (0.24) | F=0.56 (0.41), p=0.74 (0.2) |
| **Entire Sample** | **CBCL** | aggressive | F=6.96 (2.75), p=0.01 (0.01) | F=4.92 (1.79), p=0.01 (0.02) | F=0.89 (0.63), p=0.6 (0.22) |
|  |  | anxious depressed | F=0.88 (1.16), p=0.62 (0.45) | F=1.86 (0.62), p=0.16 (0.09) | F=2.12 (0.06), p=0.27 (0.06) |
|  |  | attention problems | F=11.45 (1.63), p=0 (0) | F=9.86 (1.68), p=0 (0) | F=5.07 (1.3), p=0.02 (0.02) |
|  |  | externalizing | F=9.99 (3.9), p=0 (0) | F=6.28 (2.34), p=0.01 (0.01) | F=1.91 (0.88), p=0.3 (0.15) |
|  |  | internalizing | F=0.61 (0.56), p=0.67 (0.32) | F=1.92 (0.16), p=0.15 (0.03) | F=0.79 (0.44), p=0.6 (0.18) |
|  |  | rule breaking | F=14.75 (5.65), p=0 (0) | F=8.14 (2.04), p=0 (0) | F=5.46 (1.42), p=0.01 (0.02) |
|  |  | social problems | F=6.96 (1.85), p=0 (0) | F=8.73 (3.45), p=0 (0) | F=1.09 (0.74), p=0.54 (0.24) |
|  |  | somatic complaints | F=1.15 (0.35), p=0.41 (0.16) | F=1.41 (0.58), p=0.29 (0.21) | F=1.56 (0.25), p=0.36 (0.06) |
|  |  | thought problems | F=2.8 (0.59), p=0.07 (0.05) | F=4.32 (0.69), p=0.01 (0.01) | F=1.58 (0.92), p=0.41 (0.28) |
|  |  | total problems | F=6.53 (2.05), p=0 (0.01) | F=8.14 (2.55), p=0 (0) | F=0.89 (0.58), p=0.58 (0.26) |
|  |  | withdrawn depressed | F=3.49 (2.1), p=0.13 (0.21) | F=3.45 (0.81), p=0.03 (0.03) | F=1 (0.46), p=0.53 (0.15) |
|  | **Cognitive Functioning** | common-EF-(CFA) | F=44.19 (4.33), p=0 (0) | F=30.42 (4.89), p=0 (0) | F=15.09 (2.11), p=0 (0) |
|  |  | cognitive-aptitude-(CFA) | F=72.43 (12.35), p=0 (0) | F=47.04 (4.99), p=0 (0) | F=27.2 (2.27), p=0 (0) |
|  |  | updating specific-(CFA) | F=47.42 (10.7), p=0 (0) | F=28.69 (1.67), p=0 (0) | F=21.15 (1.82), p=0 (0) |
|  |  | LMT | F=23.72 (3.69), p=0 (0) | F=14.97 (1.34), p=0 (0) | F=8.05 (3.43), p=0 (0) |
|  |  | RAVLT | F=23.54 (3.61), p=0 (0) | F=15.6 (0.73), p=0 (0) | F=8.41 (2.21), p=0 (0) |
|  |  | general-capability-(BPPCA) | F=67.07 (9.62), p=0 (0) | F=39.12 (3.14), p=0 (0) | F=25.92 (2.77), p=0 (0) |
|  |  | executive-capability-(BPPCA) | F=10.18 (2.71), p=0 (0) | F=7.15 (1.5), p=0 (0) | F=3.76 (1.05), p=0.02 (0.03) |
|  |  | learning/memory-(BPPCA) | F=31.99 (5.61), p=0 (0) | F=19.09 (0.28), p=0 (0) | F=12.18 (1.3), p=0 (0) |
|  | **Stroop** | angry acc eq | F=12.39 (3.46), p=0 (0) | F=6.02 (1.8), p=0 (0) | F=7.06 (1.79), p=0 (0) |
|  |  | happy acc eq | F=10.23 (1.67), p=0 (0) | F=6.17 (1.86), p=0 (0) | F=5.89 (2.82), p=0.01 (0.01) |
|  |  | stroop interference | F=6.85 (3.36), p=0 (0.01) | F=3.97 (0.86), p=0.01 (0.01) | F=2.42 (0.97), p=0.11 (0.14) |
|  | **UPPS** | negative urgency | F=4.15 (0.82), p=0.02 (0.03) | F=2.84 (1.34), p=0.12 (0.19) | F=4.07 (1.66), p=0.04 (0.05) |
|  |  | perseverance | F=3.73 (0.77), p=0.03 (0.03) | F=5.69 (0.88), p=0 (0) | F=2.57 (1.41), p=0.16 (0.13) |
|  |  | positive urgency | F=7.66 (2.1), p=0 (0) | F=6.89 (2.31), p=0 (0) | F=5.01 (2.47), p=0.04 (0.05) |
|  |  | premeditation | F=2.75 (1.12), p=0.09 (0.12) | F=6.11 (0.86), p=0 (0) | F=1.55 (0.8), p=0.3 (0.26) |
|  |  | sensation seeking | F=1.61 (0.54), p=0.22 (0.13) | F=1.26 (1.07), p=0.42 (0.49) | F=1.3 (1.53), p=0.51 (0.41) |
| **Supplemental Table 7.**  Bootstrapped ANOVA difference outputs across various measures of cognitive functioning and mental health by Sample. Displayed values include the mean FDR F and p-values, with standard deviations (SD) provided in parentheses. | | | | | |

| **Supplement Table 8** | | | | | | | | | | |
| --- | --- | --- | --- | --- | --- | --- | --- | --- | --- | --- |
| **Measure** | **Sample** | **10%** | **20%** | **30%** | **40%** | **50%** | **60%** | **70%** | **80%** | **90%** |
| **To Original Sample Max Correlations** | Sample-1 | 0.864 (0.17) | 0.931 (0.105) | 0.945 (0.097) | 0.971 (0.062) | 0.985 (0.019) | 0.988 (0.017) | 0.994 (0.01) | 0.995 (0.01) | 0.998 (0.004) |
|  | Sample-2 | 0.861 (0.17) | 0.923 (0.128) | 0.961 (0.07) | 0.978 (0.05) | 0.984 (0.035) | 0.992 (0.01) | 0.995 (0.005) | 0.997 (0.003) | 0.999 (0.001) |
| **Across Sample Max Correlations** | Sample-1 | 0.818 (0.157) | 0.9 (0.106) | 0.921 (0.103) | 0.954 (0.073) | 0.974 (0.028) | 0.981 (0.021) | 0.989 (0.014) | 0.991 (0.015) | 0.997 (0.006) |
|  | Sample-2 | 0.807 (0.179) | 0.883 (0.138) | 0.936 (0.087) | 0.965 (0.048) | 0.973 (0.041) | 0.985 (0.017) | 0.991 (0.009) | 0.994 (0.005) | 0.997 (0.003) |
| **Modularity (Q)** | Sample-1 | 0.449 (0.028) | 0.486 (0.027) | 0.51 (0.033) | 0.52 (0.028) | 0.538 (0.026) | 0.549 (0.022) | 0.56 (0.019) | 0.566 (0.014) | 0.571 (0.01) |
|  | Sample-2 | 0.449 (0.029) | 0.488 (0.028) | 0.517 (0.034) | 0.541 (0.023) | 0.556 (0.024) | 0.569 (0.017) | 0.579 (0.016) | 0.587 (0.011) | 0.595 (0.008) |
| **ARI** | Sample-1 | 0.588 (0.104) | 0.671 (0.086) | 0.71 (0.095) | 0.758 (0.069) | 0.806 (0.052) | 0.833 (0.049) | 0.872 (0.032) | 0.893 (0.036) | 0.933 (0.022) |
|  | Sample-2 | 0.608 (0.095) | 0.691 (0.088) | 0.762 (0.071) | 0.807 (0.044) | 0.836 (0.046) | 0.867 (0.028) | 0.893 (0.025) | 0.914 (0.017) | 0.942 (0.013) |
| **Supplemental Table 8.** Illustrates the mean and standard deviations (SD) provided in parentheses across the four down-sampled metrics of interest at varying down-sampled percentages (10% to 90%) on the two independent samples. | | | | | | | | | | |
