## Supplemental Tables for "Neurodevelopmental Subtypes of Functional Brain Organization in the ABCD Study Using a Rigorous Analytic Framework": Neurodevelopmental Subtypes Supplemental Text and Figures.docx

**S1. Reproducibility and Reliability**

To investigate the reproducibility and the reliability of the RSFC subtypes and their linkages to cognitive and emotional profiles, two key strategies are employed to ensure the reproducibility, reliability, and robustness of the identified subtypes: bootstrapping (Pat et al., 2023) and split-sample down-sampling. Bootstrapped analysis of variance (ANOVA) is used to evaluate the reliability, reproducibility, and stability of subtype differences in their cognitive and emotional profiles in the dataset by resampling individuals from each subtype with replacement. This procedure allows for estimating variability, quantifying uncertainty, and calculating confidence intervals and hypothesis tests without relying on assumptions about the underlying distribution. Split-sample sub-sampling involves randomly selecting different subsets (range 10%-90%) from demographically matched split samples to systematically assess the robustness of the subtype results obtained from the original datasets. Examining the sensitivity of the results to changes in sample size and composition allows us to determine confidence in our findings, which is crucial when working with smaller datasets where small changes can potentially significantly impact the results.

Our goal in using such rigorous methods is to enhance our findings' reproducibility, reliability, and robustness. The success of our approach in investigating the reproducibility and reliability of the RSFC subtypes in our split-sample subsampling analyses will be gauged by subtype profile (mean max correlations) and individual (adjusted rand index, ARI) similarity. High mean max correlations (average > .9) to the original and subsampled subtype profiles would indicate robustness and reproducibility in our split-sample down-sampling technique. Furthermore, an upward trend in the ARI values, consistently greater than chance as the sub-sample size increases, will further validate the reliability and reproducibility of these subtypes. For our bootstrapping analyses, success will be reflected by a mean FDR-corrected p-value < .05 across all three samples (Sample-1, Sample-2, and the Full Sample) for a given measure, suggesting our findings are statistically significant and not attributable to chance. We predict that through these rigorous methodologies, our findings will reveal robust reproducibility and reliability across the subtypes, allowing us to determine which cognitive and emotional measures robustly differ across them. These analyses will allow us to substantiate the RSFC subtypes' potential as effective or ineffective neuro-markers of cognitive functioning and emotional processing in these late school-age children.

**Supplemental Figure 1**

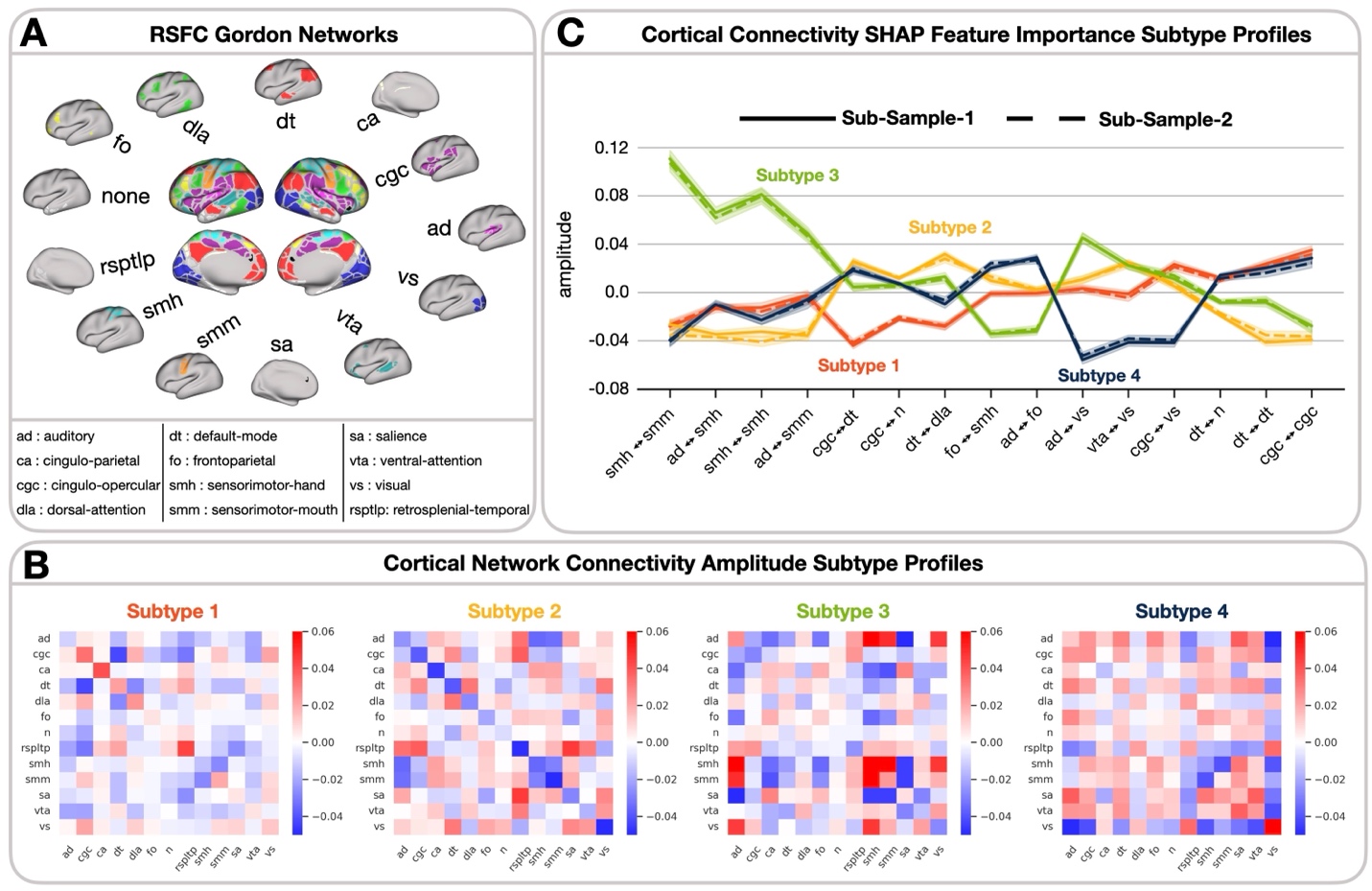

**Supplemental Figure 1.** Resting State Functional Connectivity (RSFC) Amplitude Subtype Profiles. **A)** RSFC Gordon Networks legend for **B** and **C.** All 13 networks are displayed and labeled accordingly. The *none* network consists of cortical brain regions not included in any other established network. **B)** Connectivity amplitude is denoted by blue for negative and red for positive. Given the high reproducibility across Sample-1 and Sample-2 subtypes, both samples were combined and shown for network connectivity display. are shown sequentially from left to right (Subtype-1, Subtype-2, Subtype-3, Subtype-4). **C)** Subtype profiles represent the mean age and sex-corrected functional connectivity amplitude among the top 15 cortical RSFC identified by SHAP feature importances for classifying each RSFC Subtype. Lines are colored by Subtype association (Subtype-1, orange; Subtype-2, yellow; Subtype-3, green; Subtype-4, blue), and differentiated by sample (Sample-1, straight, Sample-2; dashed).

**Supplemental Figure 2**

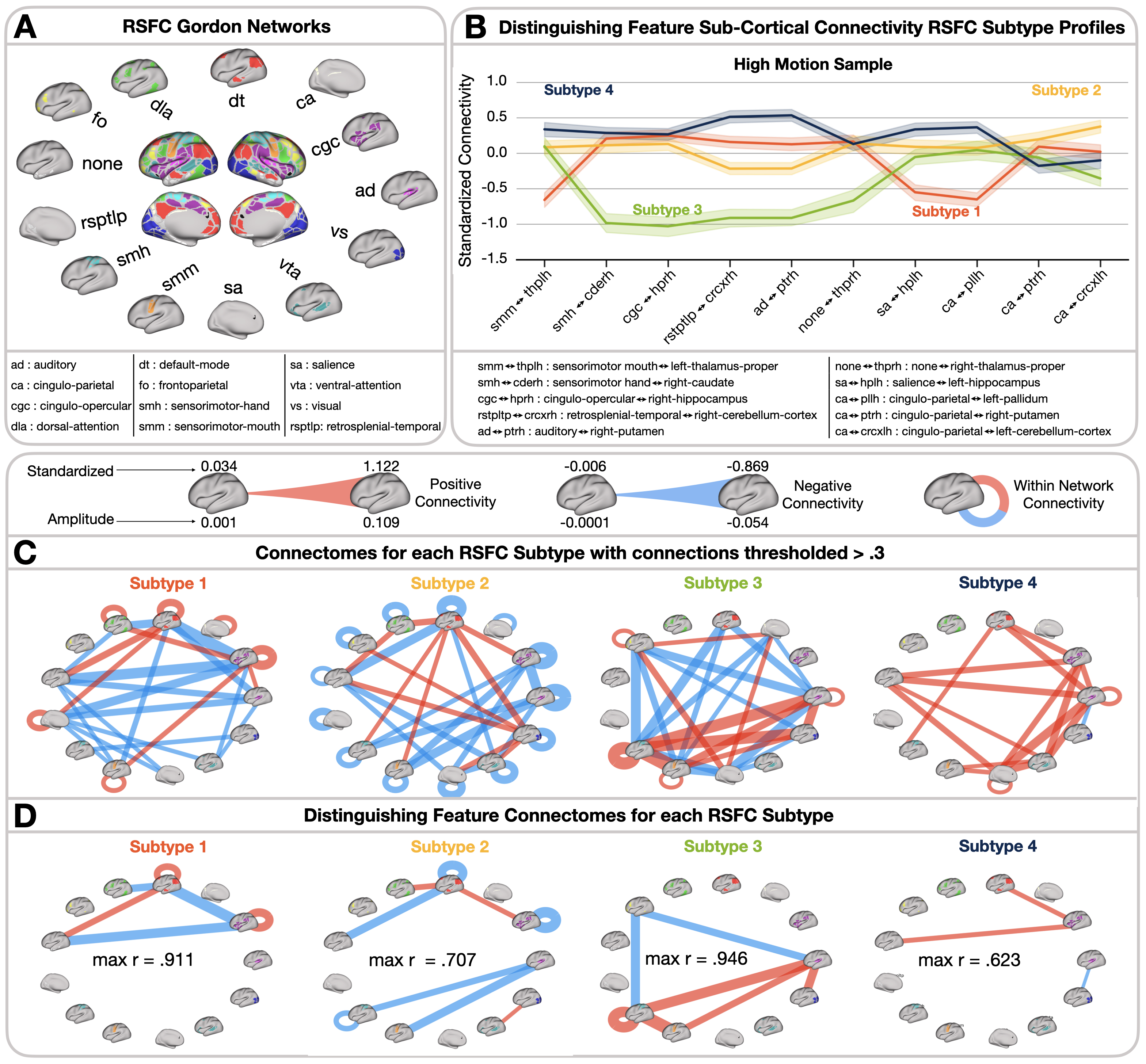

**Supplemental Figure 2.** “High Motion” Sample Resting State Functional Connectivity (RSFC) Subtype Profiles. **A)** RSFC Gordon Networks legend for **C** and **D.** All 13 networks are displayed and labeled accordingly. **B)** Subtype profiles represent the mean standardized functional connectivity among the top 10 sub-cortical regions identified by SHAP feature importances for classifying each subtype. Lines are colored by Subtype association (Subtype-1, orange; Subtype-2, yellow; Subtype-3, green; Subtype-4, blue) and differentiated by sample (Sub-Sample-1, straight, Sub-Sample-2; dashed). Full sub-cortical ROI and cortical network names are displayed below the profiles. For both **C** and **D,** the line thickness represents the connectivity strength. Connectivity directionality is denoted by blue for negative and red for positive. Self-loops characterize within-network connectivity. Given the high reproducibility across Sub-Sample-1 and Sub-Sample-2 subtypes, the Full-Sample subtypes are displayed from left to right. **C)** Subtype profiles represent the functional connectivity among cortical regions based on a connectivity threshold of 0.3 for each RSFC Subtype. **D)** Subtype profiles represent the functional connectivity among the top 15 cortical regions identified by SHAP feature importances for classifying each RSFC Subtype.

**Supplemental Figure 3**

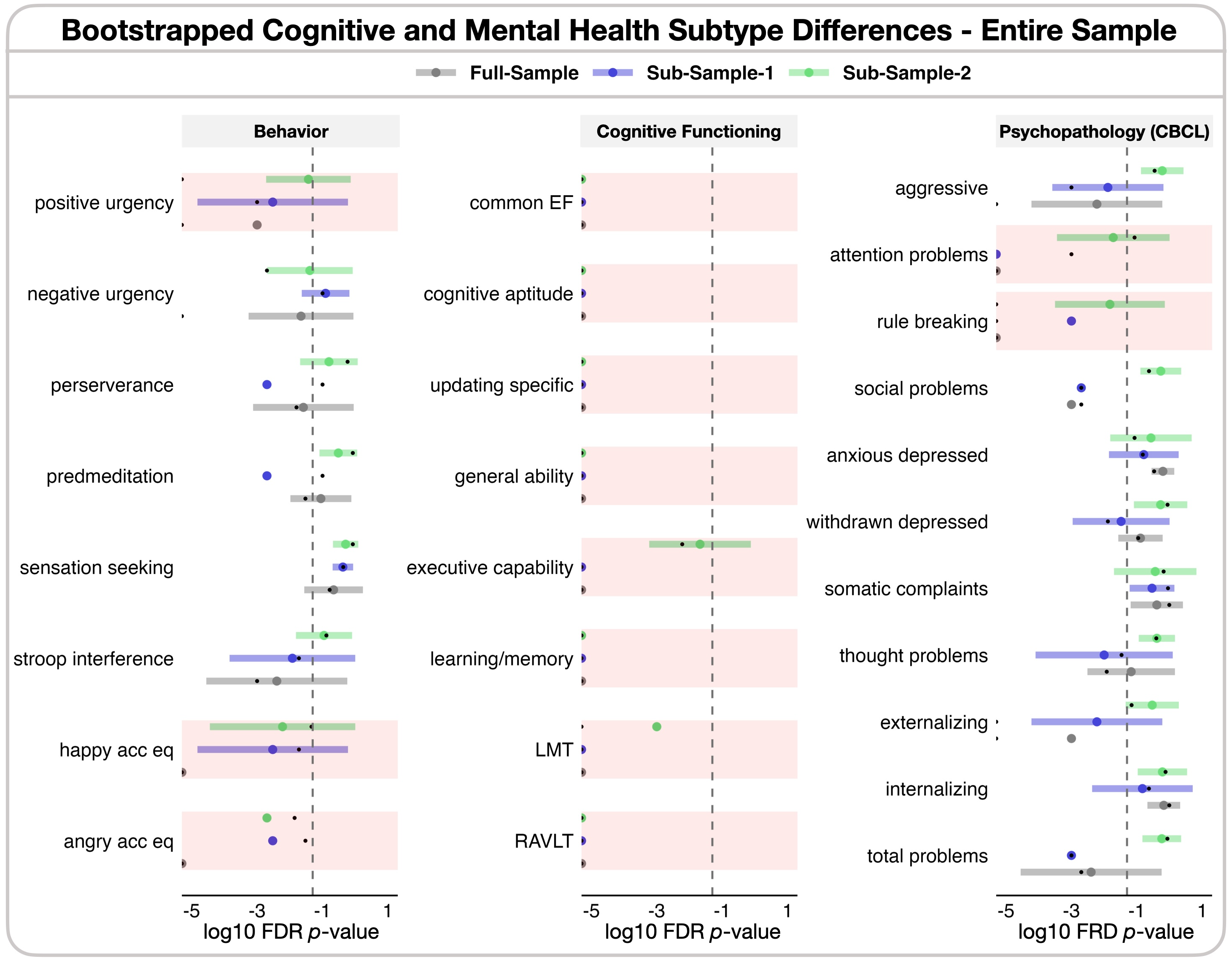

**Supplemental Figure 3.** “Entire Sample” Bootstrapped ANOVAs by Sample by Phenotypic Measure. Bootstrapped ANOVAs, log10 FDR corrected P-values from each iteration are evaluated by sample, where black dots indicate the original p values from the non-bootstrapped samples, colored dots and CI's are colored by sample (Full Sample, grey; Sample-1, blue; Sample-2; green) and represent the mean and range of p-values for each sample across the 1000 bootstrapped ANOVAs, and shaded bars in red indicate the mean of all three samples are < .05.
